# Viral infection induces oxylipin chemical signaling at the end of a summer upwelling bloom: implications for carbon cycling

**DOI:** 10.1101/2025.05.18.654734

**Authors:** Bethanie Edwards, Kimberlee Thamatrakoln, Chana F. Kranzler, Justin Ossolinski, Helen Fredricks, Matthew D. Johnson, Jeffrey W. Krause, Kay D. Bidle, Benjamin A.S. Van Mooy

## Abstract

Diatoms are large phytoplankton that form the base of the marine food web and often bloom first when nutrients are injected into the surface ocean through upwelling or deep ocean mixing^1,2^. Diatoms contribute 20% of global photosynthesis^3^ while disproportionately representing 40% of carbon export^4^, with most export occurring along the continental margins^5^. Oxylipin chemical signaling by diatoms has been extensively studied in the Mediterranean Sea where oxylipins are linked to grazing with subsequent insidious effects on copepod reproduction^6–13^. Culture studies with diatoms have shown that stress, growth phase, and viral infection also impact oxylipin production^14–16^. This study provides insight into the role of oxylipins as biomarkers and chemical signals during viral infection of diatoms in natural communities. Biomarkers for lysis and senescence were identified in laboratory experiments and observed at elevated concentrations in meta-lipidomes collected in the California Coastal Ecosystem (CCE) where diatoms had recently been lysed by viruses^17^. Deck-board incubations with natural communities show that oxylipins stimulate sinking particle-attached and surface-ocean microbes in a dose and community-dependent manner, while inhibiting microzooplankton grazing and phytoplankton growth rates. Carbon export was two times higher at the Post-lytic site than elsewhere along the transect consistent with the viral shuttle, whereby viruses facilitate carbon export. We previously reported enhanced enzymatic activity at the Post-lytic site^17^, suggestive of the viral shunt, whereby carbon is remineralized or attenuated into non-sinking dissolved organic matter. Here we layer geochemical evidence to show that lysis of oxylipin producing diatoms amplified the vertical flux of carbon from the surface ocean even in the presence of viral shunt processes. The remineralization length scale and community composition have been hypothesized as controls on shunt vs. shuttle^18–20;^ our analysis provides another example of how community interactions may toggle a system between favoring shunt or shuttle.

## Introduction

In 1999, Miralto *et al*. linked chemical signals produced by diatoms to decreased copepod egg viability during spring blooms in the Adriatic Sea (Northern Mediterranean)^6^. These chemical signals, polyunsaturated aldehydes (PUAs; Fig 1A), belong to the oxylipin subclass of lipids and resemble grazing defense molecules produced by higher plants. Subsequent research using diatom cultures explored the biosynthesis pathways of PUAs and other non-volatile oxylipins (NVOs)^14,15,21,22^, which rely on enzymatic oxidation of polyunsaturated fatty acids, specifically C22:6, C20:5, C20:4, C16:4, and C16:3. The production of oxylipins in diatoms is generally tied to stress with studies showing relationships between enhanced oxylipin concentrations and nutrient stress, copepod grazing, cell disruption, microzooplankton grazing, membrane permeability, virus infection and growth phase^8,10,12,16,23–26^.

**Figure 1.**
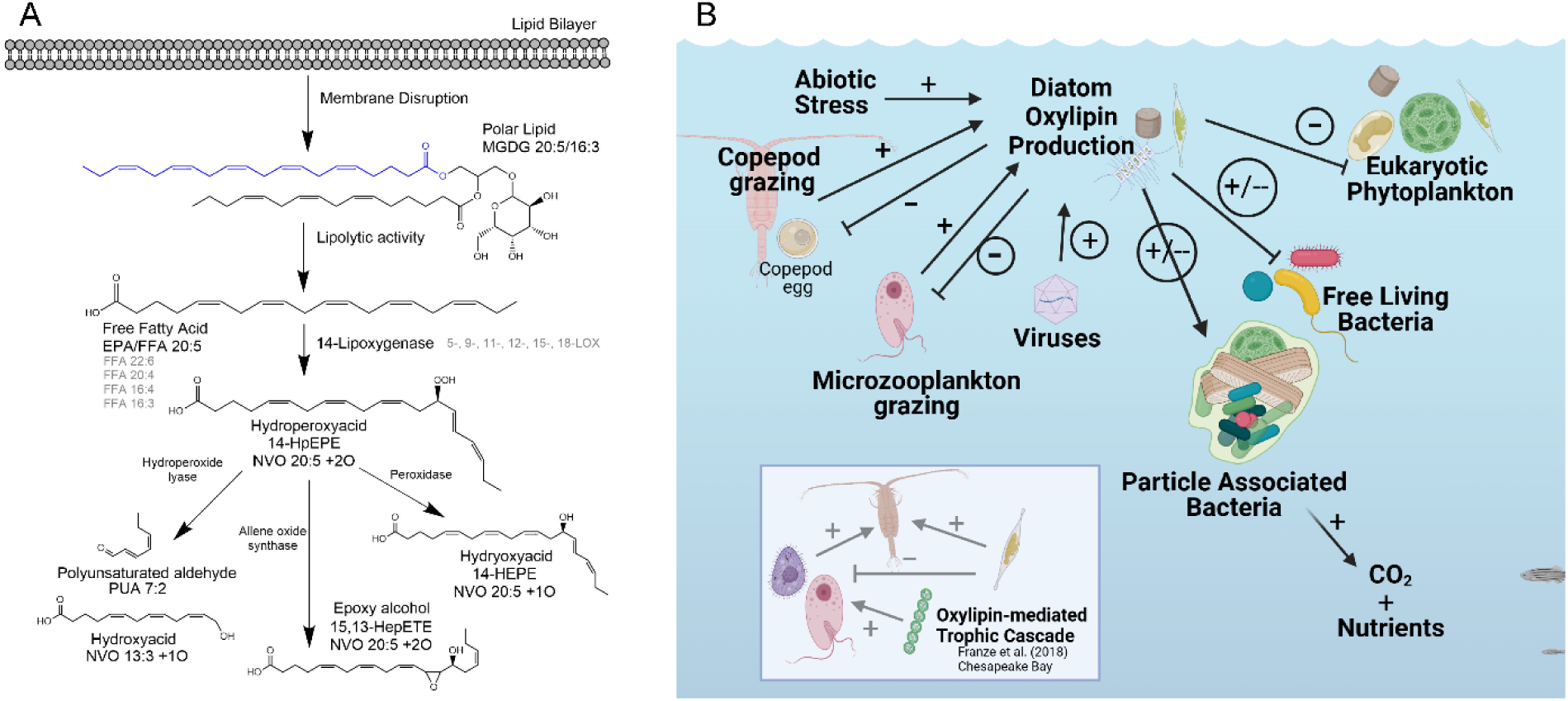
Diatoms in the ocean biosynthesize oxylipins from membrane lipids, which act as chemical signals impacting growth, behavior, cell fate, and biogeochemistry. **A)** Oxylipin biosynthesis pathway proposed for the 14-lipoxygenase enzyme acting on eicosapentaenoic acid which was lipolytically cleaved from the membrane lipid, MGDG 20:5/16:3 (monogalactosyldiacylglycerol). A hydroperoxy acid is the first compound produced, 14-hydroperoxy eicosapentaenoic acid (14-HpEPE). Enzymes downstream of LOX act on the hydroperoxy acid to produce a plethora of oxylipin compounds. Here we include other common lipoxygenases which add a hydroperoxy group (-OOH) at the specified carbon position along the fatty acid chain. **B)** Assessments of oxylipin bioactivity in surface ocean ecosystems based on culture amendment experiments and deck-board incubations^6,7,12–16,26–28,31,32,35,40,41^. Arrows denote the directionality of response and +/- denotes stimulation or inhibition, respectively (depending on concentration). Circled responses indicate interactions explored in this study. Inset shows the trophic cascade mediated by PUAs in a natural population from the Chesapeake Bay, where oxylipins decreased grazing of microzooplankton like dinoflagellates (pink flagellate) and ciliates (purple ciliated cell) on diatoms but increased grazing on cyanobacteria (green chain-forming cells) and increased copepod grazing on both diatoms and ciliates^31^.

Oxylipin amendment experiments with cultured isolates and natural communities^27–35^ (summarized in Fig. 1B) reveal a variety of cellular and ecosystem impacts. Oxylipins are largely deleterious, causing teratogenic effects on the offspring of metazoan grazers^6,7,9,10,36^, deterring microzooplankton grazing by microzooplankton^31–33,37^, and stunting the growth of eukaryotic phytoplankton competitors^27,31^. Specifically, PUAs are also a fundamental component of a nitric oxide-based, stress surveillance system whereby diatoms sense and respond to biotic and abiotic stressors, which can lead to the activation of programmed cell death at threshold concentrations^29^. At sub-lethal concentrations, PUAs can desensitize diatom cells to physiological stress and fortify cellular responses for growth and survival^29^. Lastly, PUAs have been found to be stimulatory to particle-associated bacteria and diatom epibionts in sublethal doses (1-10 µM in the Atlantic; 13-18 µM in culture), while free-living bacterial isolates tend to be inhibited at concentrations as low as 3 µM^28,35^. PUA concentrations high enough to deter microzooplankton grazing and stimulate particle-associated communities have been observed in the water column in the Mediterranean Sea^7,23,25,27,38^, within sinking particles in the Atlantic^35^, and within suspended particles in the Pearl River Estuary, China^39^ (Table S1; Fig. S1).

We previously presented a detailed comparison of the suite of oxylipins produced by three different *Chaetoceros* diatom host-virus systems, highlighting the unique oxylipins produced by each host-virus pair, and focusing on compounds with known allelopathic effects on meso- and microzooplankton grazers^16^. *Chaetoceros* species are abundant in natural diatom communities in the CCE and the most abundant diatom in the global TARA Oceans dataset^17,42–45^. Virus infection is an important loss term for phytoplankton in the marine environment with mortality rates on par with grazing across the North Atlantic and Southern Ocean^46,47^. Given lipids have been used as diagnostic biomarkers for viral infection of the coccolithophore, *Gephyrocapsa huxleyi*^48^, including in natural populations across the North Atlantic^49^, we sought to 1) explore oxylipins as biomarkers of viral infection in diatoms and 2) test the ecological relevance of oxylipin chemical signaling during infection. We took a two-pronged approach, using a laboratory model system with three diatom host-virus pairs^16,50–52^ and analysis of meta-lipidomic data from diatom assemblages in the CCE.

Three phytoplankton patches with natural diatom populations were found to be in different stages of the bloom lifecycle and viral infection (Active Infection, Post-lytic, and Non-blooming) in a 2013 field study along the CCE and established a link between viral infection and silicon limitation^17^. Our analyses showed that dissolved oxylipins in the CCE meta-lipidome were more abundant at a Post-lytic sites than at Active Infection sites or a Non-blooming sites dominated by dinoflagellates. Five of the oxylipin biomarkers from control, laboratory infection experiments were more abundant *in situ* at the Post-lytic site coinciding with higher bacterial ectoenzyme activity^17^ and higher Si-stress^17,43^ compared to the Active Infection site. Deck-board incubations exposing surface and sinking particle-associated microbial communities to exogenous oxylipins suggest that they impact grazing, growth, and organic matter cycling by marine microbes. At the same time, sediment traps and satellite data confirm the higher oxylipin post-lytic siters also had higher magnitude and efficiency of carbon export.

## Results and Discussion

### Viral Infection and mortality biomarkers identified in culture experiments

Two diatom species (*Chaetoceros tenuissimus* strain 2-10 *and C. socialis* strain L-4) served as hosts in laboratory infection experiments with three taxonomically distinct viruses: a single-stranded DNA (ssDNA) virus (CtenDNAV) and a single-stranded RNA (ssRNA) virus (CtenRNAV), both infecting *C. tenuissimus;* and a ssRNA virus (CsfrRNAV), infecting *C. socialis.* Control and infected treatments were sampled for lipidomics and cell concentration over 3-4 days post infection (dpi; Fig. 2A, S2). Samples were categorized based on the growth phase (Exponential, Stationary, and Decline) and dissolved lipidomes were annotated as described in Edwards et al.^16^. We evaluated the general lipidomic response across the three host-virus pairs to elucidate viral infection and senescence responses shared across the genus *Chaetoceros*.

**Figure 2.**
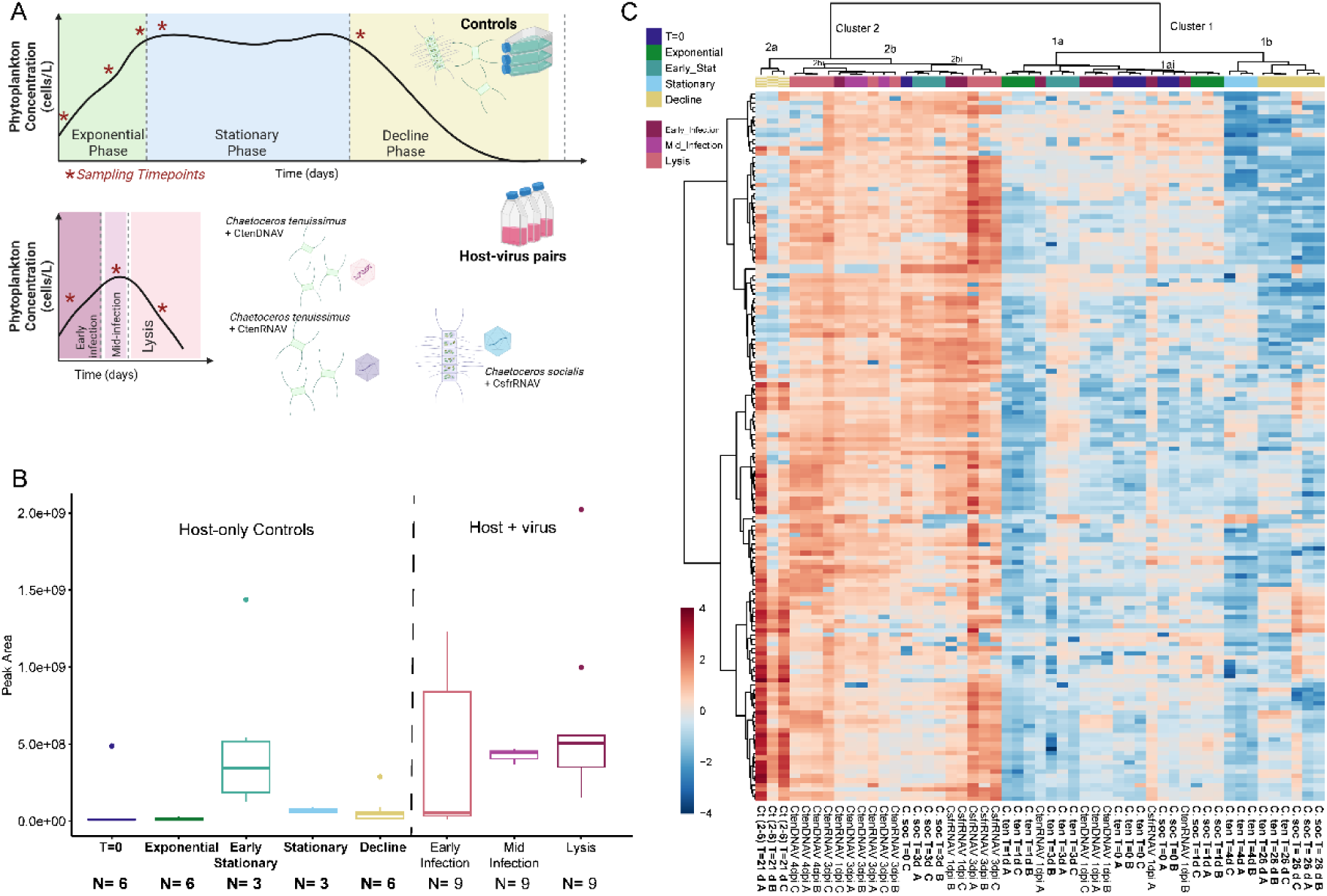
Virus infection of diatoms leads to the release of oxylipins and free fatty acids into the dissolved organic matter pool. **A)** Experimental design showing a representative growth curve of the host-only control and representative time course of infection for three diatom host-virus pairs: The approximate sampling timepoints are noted with red asterisks. The actual growth curve of each triplicate treatment can be found in Fig. S2. **B**) Box and whisker plots of the average peak area of oxylipins and free fatty acids in the dissolved lipidome for host-only controls and the stage of infection averaged across all three viruses. The edges of the box indicate the 1^st^ to 3^rd^ quartile range; error bars represent the standard deviation with outliers noted by dots. **C)** Heatmap of the oxylipidome (Oxylipins + FFA) from the *Chaetoceros* viral infection experiments showing the relative abundance of each compound relative to its mean concentration across all samples with red indicating a relative increase and blue a relative decrease, reported as a log fold change. Colored bars above the heatmap denote the growth phase and infection stage as indicated. The samples on the X-axis and compounds in the Y-axis are ordered based on the dendrogram created using Ward clustering. Important clusters of samples that are discussed in the text are labeled on the dendrogram. Timepoints as labeled by color at the top of the heatmap and individual samples are labeled at the bottom of the heatmap. Controls are labeled with the host name, day sampled after transfer, and triplicate identifier, A, B, or C. Infected samples are labeled with the virus, days post infection (dpi), and triplicate identifier, A, B, or C.

The dissolved lipidome (<0.2 μm) represents small and often oxidized lipids that are more soluble than hydrophobic storage lipids or polar membrane lipids. High Resolution Accurate Mass-Mass Spectrometry (HRAM-MS) paired with reverse phase ultra-high performance liquid chromatography analysis was used to analyze the dissolved lipid extracts and reference standards (Table S2). To address specific questions about oxylipin abundance and diversity, we focused on the 156 compounds confidently annotated as free fatty acid (FFA) precursors of oxylipins and NVOs. Equal weight was given to high and low abundance compounds by mean-centering and log-transforming the peak area data after normalizing to the volume filtered and the recovery of an internal standard (Supplemental File 1). A comparative -omics scheme^53,54^ was applied to the resulting dissolved oxylipin profiles using hierarchical clustering analysis and multivariate analysis to determine significant biomarkers of viral infection, stress, and death.

*Chaetoceros* produced the highest absolute abundance of FFAs and NVOs (*i.e.*, non-PUA oxylipins) during the later stages of viral infection (mid-infection and lysis) and in early stationary phase in controls (Fig. 2B-C). Hierarchical clustering analysis split the data into two clusters revealing distinct lipidome structures between the cells lysing due to viral infection (Cluster 2) and the aging, control treatments (Cluster 1; Fig. 2C). Within Cluster 2, 15 of 22 samples are infected, and within Cluster 2bi, all 10 samples are from mid-infection and lysis of *C. tenuissimus*. This suggests that *C. tenuissimus* lipidomes from virocells are more similar to one another than to other species infected with similar viruses. Triplicates from two controls grouped with the T= 3 and 4 dpi virus samples, stationary phase *C. socialis* and declining *C. tenuissimus* from a different strain (2-6) than the CtenDNAV and CtenRNAV host (2-10). The C. *tenuissimus* host lipidome (2-10) grouped with the declining *C. socialis* host control in Cluster 1. Although *C. tenuissimus (2-6)* was not used as a host in our virus infection experiments, it was included in our metanalysis as an outgroup and a unique lipidomic signature, forming its own subgroup within Cluster 2 (Fig. 2C, Cluster 2a.). *C. socialis* showed elevated oxylipins in stationary phase with an overall lipidome most similar to CsfrRNAV infected cells (Fig. 2C, Cluster 2bii samples labeled C. soc T=3 d).

Cluster 1 was enriched with control samples (23 of 29 samples). The declining control samples from *C. tenuissimus* (2-10) and *C. socialis* clustered with stationary phase *C. tenuissimus* (2-10), and showed lower relative abundances of most oxylipins observed in the experimental dissolved meta-lipidome (Fig. 2C-Cluster 1b). The only virus infected treatments in Cluster 1 are early infection samples (T= 1 dpi; 6 of 29 samples).

The observed strain-level variability in the relative abundance and chemical diversity of oxylipins produced was consistent with Wichard *et al.*, which surveyed PUA production in 50 strains of phytoplankton (primarily diatoms)^16,22^. Ribalet *et al.* had previously established that the diatoms, *Skeletonema* and *Thalassiosira,* increase the production of free fatty acid and PUAs as they reach stationary phase and even more so under nutrient stress, particularly Si-limitation^14,15^. Our method was not optimized for PUA quantification, but rather for the pool of less volatile compounds adjacent and upstream of PUA production (Fig. 1A), showing plasticity in the range of non-volatile oxylipins produced by *Chaetoceros* diatoms under different growth conditions.

Sparse Partial Least Squares - Discriminant Analysis (sPLS-DA) was used to distill the dataset down to the twenty most distinctive features that best differentiate time-points and treatments from each other, in hopes of identifying discrete biomarkers for viral infection or general cell death (Fig. 3). Compounds that increased over the course of infection and were most abundant during lysis were assigned to Component 1 (Fig. 3B). The strongest viral biomarkers were compounds that were only abundant in the control on the transfer day (T=0) and increased with viral infection (NVO 20:2 +1O, NVO 22:4 +1O, NVO 20:3 +1O, and NVO 14:2 +1O; Fig. 3B). While the other Component 1 compounds (a different NVO 14:2 +1O isomer, NVO 22:5 +1O, NVO 16:3 +3O, two isomers of NVO 20:4 +1O, and 11,14-dihydroxy eicosapentaenoic acid) increased with infection but were present in early stationary phase as well. Compounds that were more associated with a general death response (*i.e.*, high during decline of the control and during lysis of the infected cultures) were assigned to Component 2 (Fig. 3C).

**Fig 3.**
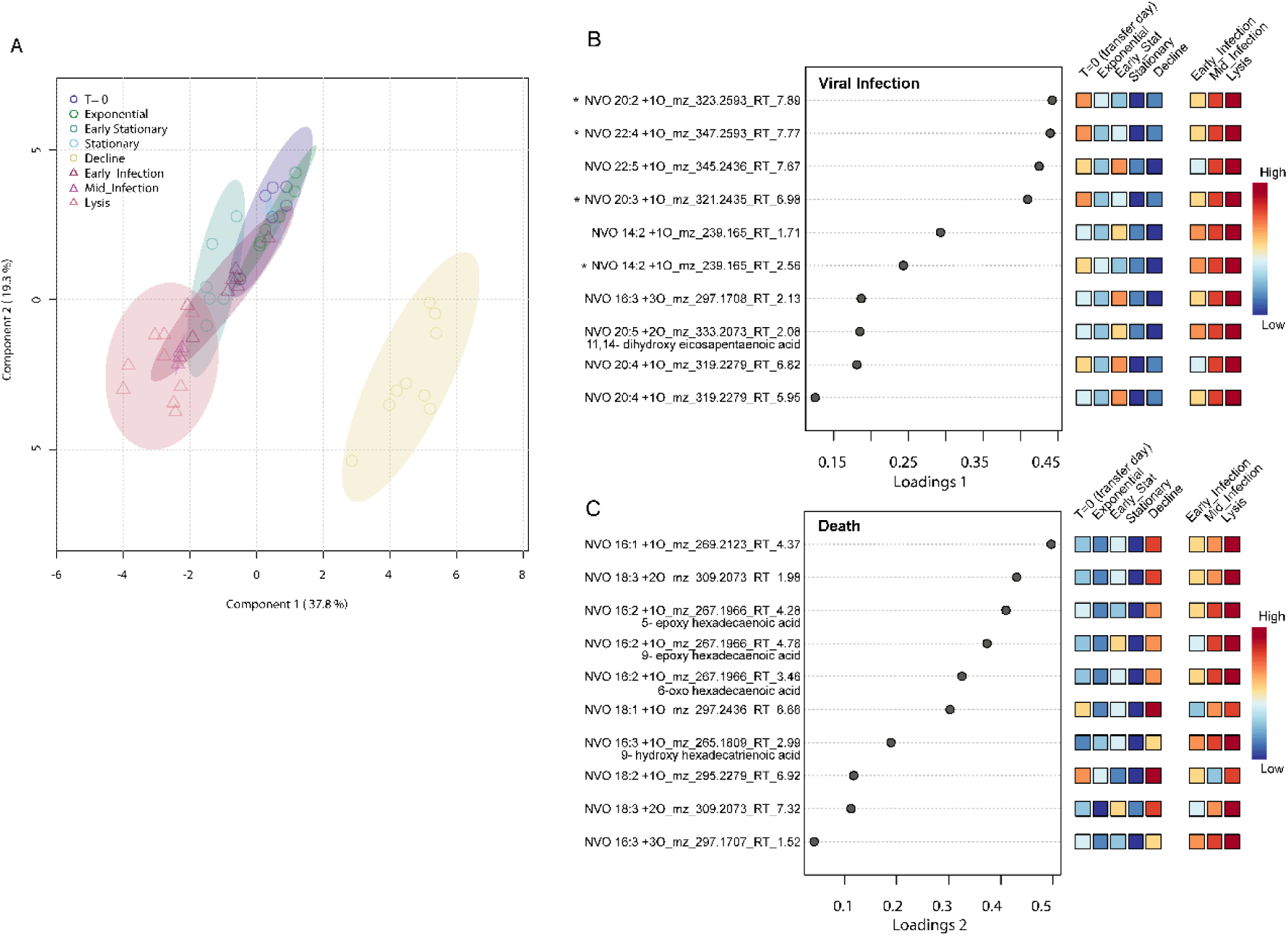
Biomarkers for virus infection and general cell death of diatoms. **A)** Loadings of the dissolved lipidome samples on Component 1 and 2 of the sparse PLSDA. Control samples are represented by a colored circle corresponding to the time of sampling (T=0, exponential, early stationary, stationary, and decline) and infected treatments are represented by pink triangles of various shades. show growth phase and viral infection lead to distinct changes in the dissolved lipidome. **B-C)** The significant compounds assigned to Components 1 and 2 by the sPLS-DA, their loadings on either component are shown as a dot-plot and the heatmap shows the average relative abundance of each compound across each timepoint. **B)** Component 1 separates the viral infection lipidomes from the decline lipidome. **C)** Component 2 captured the signal from cell death more generally.

Twelve of the twenty significant oxylipins were close analogs of known allelopathic compounds produced by diatoms that negatively influence copepod reproductive success in the Mediterranean Sea and microzooplankton grazing in the laboratory^8,10,25,33^ (Table S3). These results suggest viral infection triggers a pulse of chemical signaling via oxylipins that is distinct from the shared suite of oxylipins produced by dying, uninfected cells. This chemical signaling could impact trophic interactions and the associated biogeochemistry. We tested this hypothesis by analyzing a dissolved lipidome dataset from a field study in the California Coastal Ecosystem (CCE), where we previously characterized different stages of viral infection and conducted deck board amendment experiments with different communities using a range of oxylipin concentrations.

### Bloom decline and viral infection are linked to oxylipin production and efficient export *in situ*

#### Biogeochemical evidence of bloom decline and enhanced export

The DYEatom cruise (June 27^th^ – July 6^th^, 2013; *R/V Point Sur*) occupied nine stations within the CCE upwelling zone. The physical oceanography, phytoplankton composition, Si-stress, and infection state have been previously characterized^17,43^. Briefly, near-shore stations outside Monterey Bay were dominated by diatoms, particularly *Pseudo-nitzschia*, that were characterized as being actively infected (Active Infection) at the time of sampling. Near-shore stations by Point Reyes were also diatom-dominated but were more diverse in species composition (see Fig. 1c in Kranzler et al.^17^); they were further characterized as having undergone a recent viral-mediated mortality event (Post-lytic). Here, we present an analysis of the dissolved meta-lipidome from these two contrasting regions, along with two offshore stations that were dominated by dinoflagellates that serve as functional controls (Fig. 4, 5, S3).

**Figure 4.**
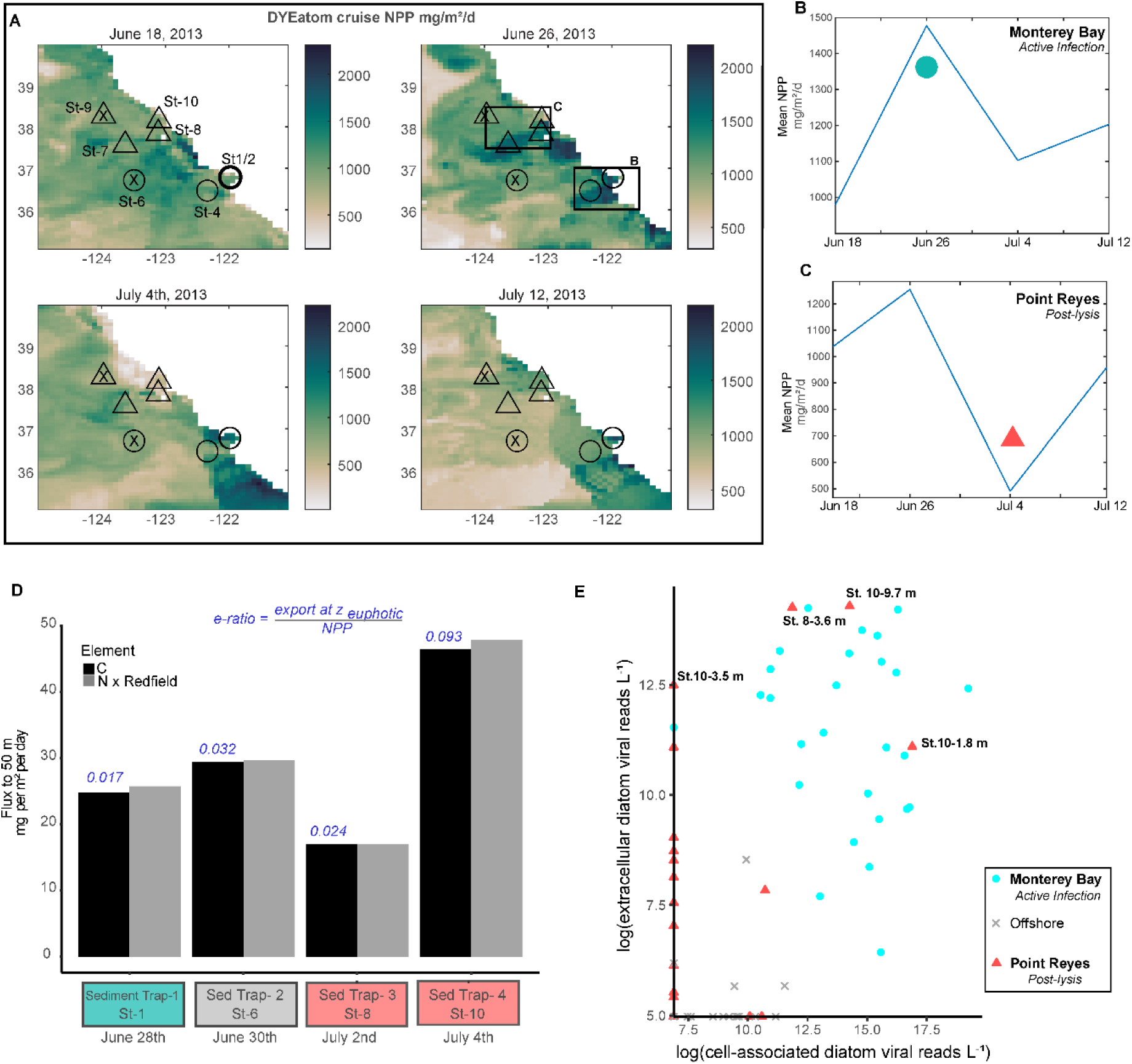
Diatom-virus dynamics, net primary productivity, and export signatures. A) A map of sampling sites during the DYEatom cruise (June 27 -July 6, 2013) and associated net primary productivity (NPP; dark green gradient) integrated over 8-day bins from June 18, 2013 to July 12^th^, 2013 using the CAFE Model^52^ of MODIS satellite data. Open circles indicate Monterey Bay stations; open triangles indicate Point Reyes stations. The stations marked with an ‘x’ indicate non-bloom site outside the upwelling filaments. Time course of the mean ocean NPP within the 1° latitude by 1° longitude rectangles drawn around **B)** Monterey Bay and **C)** Point Reyes. Filled symbols represent when Monterey Bay (cyan circle) and Point Reyes (coral triangle) were sampled. **D)** Particulate organic carbon (POC; black bars) and particulate organic nitrogen (PON; grey bars scaled to C by multiply by 6.6) flux to 50 m, as determined using sediment traps deployed at four stations over different stages of bloom decline. The blue text above each black bar indicates the export ratio calculated from the POC data and the NPP satellite estimates. **E)** Data reproduced from Kranzler *et al.* (2019)^17^ showing cell-associated diatom virus and extracellular virus copy numbers for three contigs identified as diatom RNA viruses similar to those used in culture experiments from Fig. 3^50,51^.

**Figure 5.**
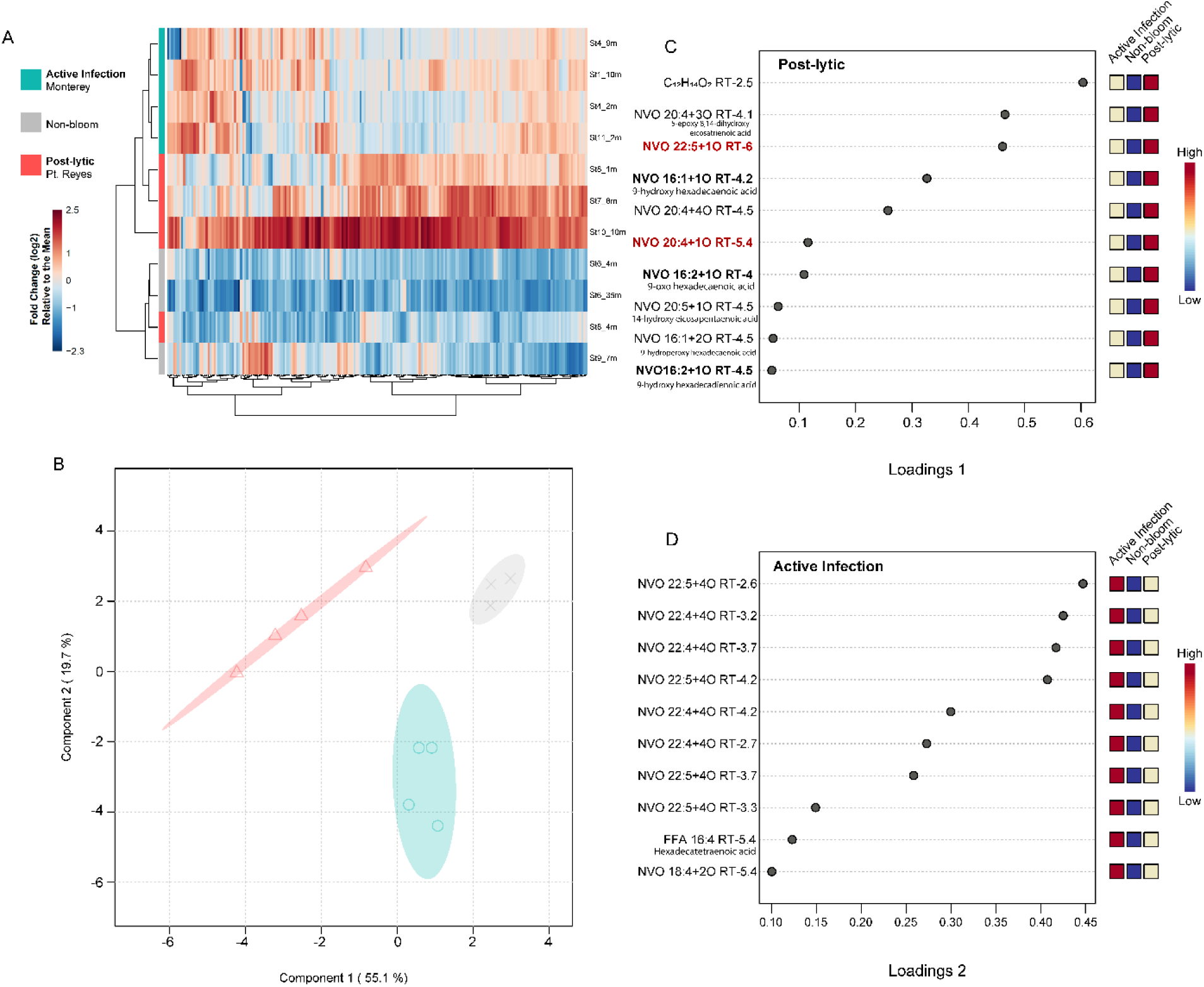
Oxylipin biomarkers of infection state from the DYEatom meta-lipidome. **A)** Heatmap showing the relative abundance of 175 compounds identified as free fatty acids and oxylipins in eleven dissolved lipidome samples from the DYEatom cruise. Samples on the X-axis (lipid compounds) and Y-axis (station and depth) are organized by similarity depicted by the Ward clustering in the dendrogram. **B)** Loading of the dissolved meta-lipidome samples on Components 1 and 2 derived from sPLS-DA for samples collected at Point Reyes (coral triangles), Monterey Bay (cyan circles), and non-bloom sites (grey x). The 95% confidence intervals (colored ellipses) for the three bloom states do not overlap. Component 1 (55.1% of the variability in the data) distinguishes the post-lytic samples from the other two categories; Component 2 (19.7% of the variability in the data) describes active infection. **C-D)** The ten compounds from the dissolved meta-lipidome used to derive **C)** Component 1 and **D)** Component 2 in the sPLS-DA. Compounds are ordered by their loading as noted by the dot-plot. Heatmaps show the average relative abundance of each compound in the dissolved meta-lipidomes from each bloom state. Compound annotations in red text are biomarkers of viral infection in *Chaetoceros* cultures (Fig. 3B), while those in bolded black text are more general cell death biomarkers (Fig. 3C).

Mean net primary productivity (NPP) in the region during our occupation was derived from satellite data and the CAFE algorithm^55^ (Fig. 4B-C). This analysis showed that we sampled an Active Infection at the Monterey Bay sites (Fig. 4E), coincident with a peak in NPP (Fig. 4B). The Post-lytic Point Reyes site had the lowest NPP at the time of sampling, consistent with a decline in productivity and diatom bloom demise. *In situ* measurements of chlorophyll, a common proxy for phytoplankton, further supported the distinction between sampling sites with concentrations between 8-18 μg L^-1^ in Monterey Bay and 3.9-7.5 μg L^-1^ at Point Reyes^43^.

Sediment traps were used to quantify carbon export out of the mixed layer (to 50 m) and calculate export efficiency as an apparent e-ratio (export at Z_euphotic_/NPP; Fig. 4D). The carbon flux in Monterey Bay was similar to the offshore, non-diatom dominated station (24.7 and 29.3 mg m^-2^ d^-1^, respectively), although the latter had a ∼2-fold higher e-ratio of 0.032 (compared to 0.017). The highest carbon flux (46 mg m^-2^ d^-1^) and highest e-ratio (0.093) were observed at the Post-lytic Point Reyes site on July 4^th^ (St. 10), suggesting export efficiency increased as viral infection and bloom declined progressed. The lowest measured carbon export flux was two days prior at a nearby station also characterized as Post-lytic St. 8), reflecting the time-resolved connection between infection dynamics and episodic carbon export^56–59^. Unlike St. 8, St. 10 still had detectable levels of intracellular virus (Fig. 4E), which may help explain differences in export. St. 10 was closer to the origin of upwelling and the populations appeared to be earlier in the lytic process while St. 8 was more offshore and further along in the lytic cycle. It is possible that St 8 was sampled after an export event had already occurred, as is evidenced by the removal of viruses from both the free-living and cell-associated fraction presumably by adsorption onto particles (a known sink for viruses from the water column^60^).

We recognize that the type of sediment trap deployed, a surface-tethered net trap, is not as quantitative as a particle interceptor trap (PIT) array or neutrally buoyant trap in determining quantitative fluxes^61^; consequently, we may be underestimating export. However, given the same net traps were used across all stations, these caveats (high precision, low accuracy) should be uniform, allowing intercomparison of export between sites. Furthermore, the highest e-value measured (0.093) notably aligns with regional estimates from Henson *et al.* derived as a function of SST^62,63^ although it is lower than an estimated global average e-ratio of 0.17^62^ and regional estimates from other models based on satellite color and food web dynamics.

#### Dissolved meta-lipidome reveals abundance of death and virus biomarkers in Post-lytic communities

Within the dissolved meta-lipidome, 175 compounds were annotated as oxylipins or free fatty acid precursors (Supplemental File 2). The Post-lytic site had a higher relative abundance of dissolved organic lipids compared to the Active Infection or non-bloom sites, especially on July 4^th^ (St 10; Fig. 5A). The Active Infection and Post-lytic lipidomes were more similar to one another than to the non-bloom lipidomes, clustering together in the sample dendrogram (Fig. 5A). sPLS-DA was used to identify the most significant features in the dissolved lipidome for differentiating between the bloom states. Component 1 represented 55.1% and Component 2 represented 19.7% of the respective variability in the data (Fig. 5B), with the former containing compounds significantly associated with the Post-lytic stage of infection (Fig. 5C) and the latter compounds more associated with Active Infection (Fig. 5D).

Three general death biomarkers and two viral lysis biomarkers from the *Chaetoceros* lab experiments were amongst the most significant compounds associated with the Post-lytic site (Fig. 5C, bold and red compounds, respectively; Table S5). The general death biomarkers 9-hydroxy hexadecadienoic acid (9-HHDE), 9-hydroxy hexadecaenoic acid (9-HHME), and 9-oxo hexadecaenoic acid (9-oxoHME) all belong to the 9-LOX pathway acting on 16:n fatty acids previously observed in *Skeletonema marinoi* and *Thalassiosira rotula* (Table S3). The abundance of the viral lysis biomarkers, 20:4 +1O and 22:5 +1O, was too low to gain positional information from ms^2^ fragmentation, but a retention time model indicates these oxylipins are hydroxy acids similar to those made by *T. rotula*, *Stephanopyxis turris*, and *Leptocylindrus* (Table S3; Supplemental File 3).

The most significant compounds in the dissolved lipidome from the Active Infection site were eight highly oxidized C22:n oxylipin isomers, the FFA and oxylipin precursor, hexadecatetraenoic acid (C16:4), and a C18:4 oxylipin (Fig 5D). While less oxidized DHA oxylipins, 22:5 +1O and 22:4 +1O, were prominent viral infection biomarkers in lab infection experiments (Fig. 3B), the highly oxidized +3O and +4O compounds associated with the Active Infection site are unique for marine systems (Fig 5D). However, C22:n oxylipins alone may not be specific enough to be indicators of viral infection. For example, C22:n biosynthesis pathways have been studied in *Leptocylindrus* diatoms showing that, in response to sonication, 15- and 18-lipoxygenases are employed to make +1O and +2O C22:n oxylipins^64^. We previously observed C22:6 +2O oxylipins produced in culture experiments feeding *Phaeodactylum tricornutum* to the dinoflagellate *Oxyrrhis marina* ^33^. Hydroxy and epoxy-hydroxy DHA derived oxylipins have been found in other natural systems, constituting approximately 20% of the total NVO pool from January to December 2017 at the LTER-MareChiara in the Mediterranean Sea where particulate oxylipin concentrations were positively correlated with diatom density^25^. The highly oxidized nature (+3O and +4O) of the C22:n oxylipins at the Active Infection site may be related to the stress-level of the community and should be further investigated as biomarkers for infection vs. +1O and +2O C22:n oxylipins, which may be more general stress biomarkers.

### Oxylipins function as chemical signaling across trophic levels in the California Current

Similar to the compounds identified in laboratory experiments, NVOs associated with Post-lytic stage of infection at Point Reyes are also known to have allelopathic effects on microzooplankton grazers in culture and copepods at sea^8,10,33^ (Table S5; Fig. 5C). An isomer of the NVO 20:4 +3O oxylipin, annotated as a 5-epoxy-8,14-dihydroxy eicosatrienoic acid, was associated with dinoflagellate grazing deterrence and chronic stress in *P. tricornutum* experiments^33^. Furthermore, the C16:1, C16:2, C20:4 and C20:5 oxylipins from the Post-lytic site have been observed in diatom blooms in the Adriatic Sea that are detrimental to copepods^8,33^.

Unlike grazers, the effects of oxylipins on eukaryotic phytoplankton and bacteria have only been widely studied using commercially available PUAs, not NVOs^23,27,28,30,34,35^. NVOs are upstream in the oxylipin biosynthetic pathway from PUAs (Fig. 1A) and tend to be toxic at lower doses than PUAs when tested against microzooplankton grazers in the laboratory^33^ and copepods at sea^8,10,13^. To assess the role of NVOs and PUAs in signaling across trophic levels, we conducted deck-board incubations of surface assemblages and sinking particle-associated microbes treated with exogenous PUAs and NVOs (Fig. 6) and assessed trends in oxylipin concentration and microbial metabolism.

**Figure 6.**
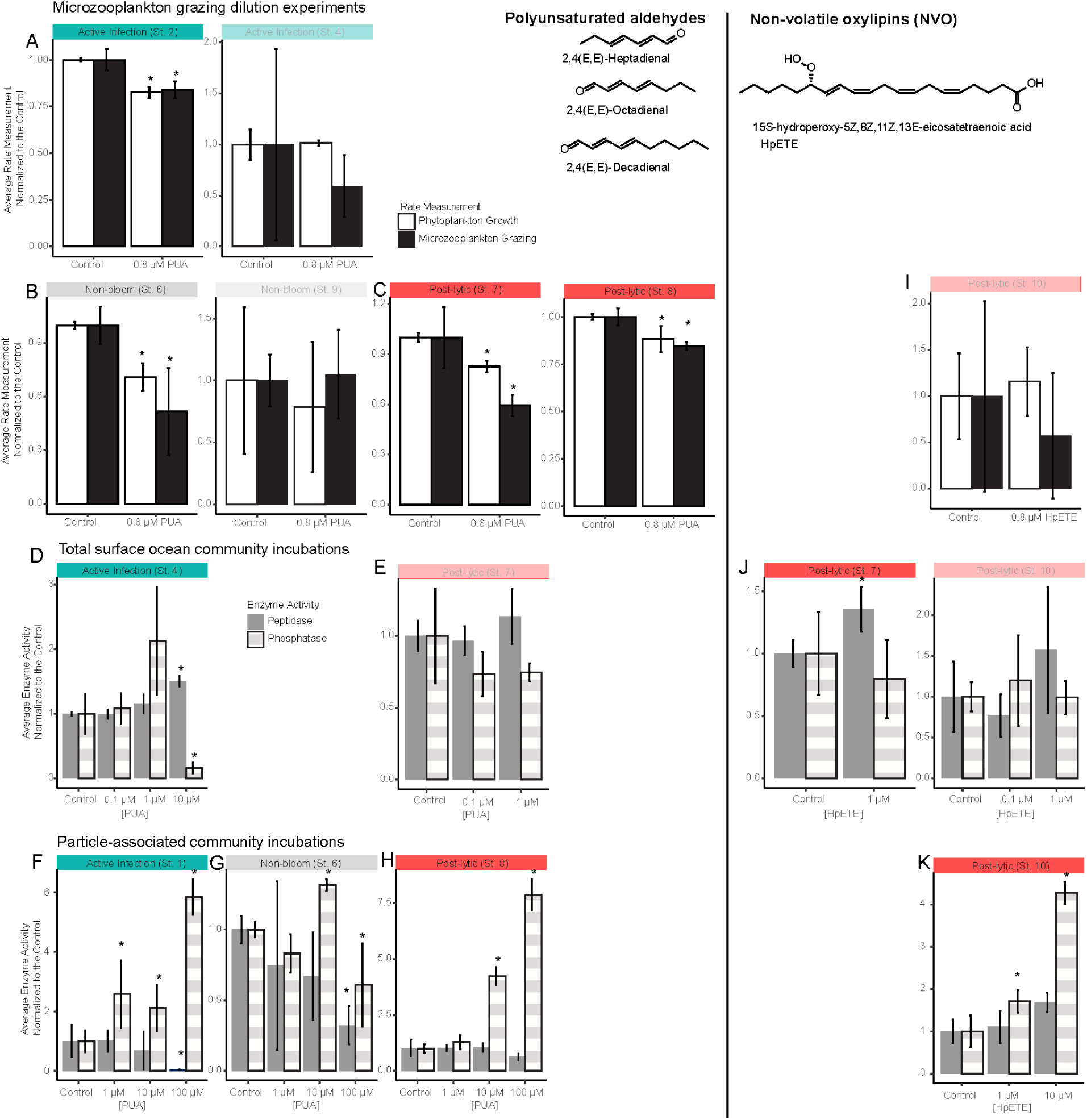
Microzooplankton grazers, phytoplankton, and both the surface and the particle associated heterotrophic microbial communities exhibit dose- and community-dependent responses to specific PUAs and non-volatile oxylipins. **A-C, I)** Fold-change in growth rate of phytoplankton (white) and grazing rate of microzooplankton (black) for control (untreated) and oxylipin amendments at seven stations. The mean and standard deviation are shown (n=3). Asterisks denote a statistical difference from the control (p < 0.05 Mann-Whitney U). Shaded panels denote statistically insignificant results. **D-E, J)** Enzymatic activity of the surface and **F-H, K)** sinking particle-associated microbial communities in response to varying concentrations of a PUA mixture **(D-H)** and the oxylipin 15-hydroxyeicosapentaenoic acid (15-HpETE; **I-K**). Alkaline phosphatase activity is in grey and white stripe and peptidase activity is in solid grey. Values are normalized to the triplicate controls. Asterisks denote a statistically significant difference from the control (p < 0.1 Krushal-Wallis with a p < 0.05 Dunn’s post-hoc).

#### Negative feedback loop between oxylipins and grazing

A mixture of PUAs—heptadienal, octadienal, and decadienal—had differential effects on natural microbial communities from the California Current. PUAs inhibited microzooplankton grazing and phytoplankton growth (Fig. 6A-C), but stimulated hydrolytic enzyme activity (peptidase and phosphatases) in surface communities and sinking particles (Fig 6D-H). A map of grazing, growth, and a proxy for oxylipin concentration (total peak area of oxylipins and FFA in the dissolved lipidome) across the study site showed the highest oxylipin concentrations co-occurred with the lowest grazing rates and the lowest phytoplankton growth rates (Fig. S4), consistent with laboratory-based interactions^33^.

The relationships between oxylipin concentration, grazing and growth across the cruise sites reflect the dose dependence of the allelopathic effects of oxylipins and the community’s threshold response to these chemical cues. Excluding the highest oxylipin concentration, a positive relationship between the oxylipin concentration proxy and phytoplankton growth was observed (Fig. S5 C). The highest oxylipin concentrations occurred at the Post-lytic Point Reyes site where the oxylipin threshold for inhibiting phytoplankton had been crossed, resulting in negative growth rates (Fig. S4 A, D).

A negative feedback loop between grazing and oxylipin production was observed, whereby grazing induces oxylipin production, which further inhibits grazing (Fig. S5 D). The lowest oxylipin concentrations were observed at stations with low grazing rates. Subsequently, a negative correlation between grazing and oxylipin concentrations was observed at the four stations with the highest oxylipin concentrations once the threshold concentration for allelopathy had been surpassed (Fig. S5 D). Several studies have documented synergy amongst various oxylipins, whereby the presence of multiple oxylipins leads to a lower effective concentration^11,13,65^. Thus, the higher diversity of lipids at the Post-lytic sites compared to the Active Infection sites (Table S6 and S7) may also contribute to the comparatively lower grazing rates (Fig. S4).

#### Dose-dependent PUA signaling in surface and sinking particle-associated microbial communities

We tested the sensitivity of surface and sinking particle-associated microbial communities to oxylipins by exposing them to a range of concentrations based on thresholds observed in the literature (Fig. S1; Materials and Methods). Depending on the bloom state and community, the response to exogenous oxylipins varied with shifting effective concentration thresholds and sign of response (negative vs. positive). At the Active Infection and Post-lytic sites, where background oxylipin concentrations were relatively high, addition of 100 μM PUAs to sinking particle-associated communities increased phosphatase activity and inhibited peptidase activity (Fig. 6F,H). The communities on particles collected from the lower-oxylipin dinoflagellate-dominated, non-bloom site (St 6) were inhibited for both enzymes after exposure to 100 μM PUA, but stimulated by a 10 μM addition (Fig 6G). Thus, the microbial communities that had prior exposure to oxylipins were more tolerant to exogenous oxylipin additions. Our findings agree with experiments conducted across the Pacific in the South China Sea showing stimulation of particle-associated communities in response to 10-100 μM PUAs^34^. The threshold of inhibition was lower in particle-associated communities from the Atlantic, where inhibition of enzyme activity, respiration, and bacteria growth was observed at 100 μM^35^ and stimulation between 1 and 10 μM PUA addition. The community dependent response to PUAs—inhibition of both peptidase and phosphatase activity on particles collected from non-blooming sites at 100 μM (Fig 6G) —is consistent with a screening of 33 bacterial isolates showing stimulation only amongst strains known to be epibionts of diatoms^22^

In surface microbial communities from the Active Infection site, peptidase activity was stimulated and phosphatase activity was inhibited at 10 μM PUA (Fig 6D). Unfortunately, we did not replicate the 10 μM PUA treatment at the other sites; so it is unclear whether non-blooming and Post-lytic surface ocean communities are more, less or equally sensitive. This oxylipin stimulation of surface ocean microbes lies in contrast to a previous study in the NW Mediterranean that found a significant decrease in single cell activity for a subset of the bacterial community in response to nM concentrations of PUAs^66^. Another study testing daily 1-2 μM PUA additions in 21-day mesocosms found no effect on microbial composition in response to PUAs alone but did see a difference in bacterial community structure with time and in the two mesocosms that ended up with different strains of *Skeletonema* blooming after nutrient addition^67^. We previously reported enhanced peptidase activity at the Post-lytic site likely reflecting the stimulatory effect of oxylipins on enzyme activity^17^. Culture studies show enhanced peptidase activity in infected *Chaetoceros* cells compared to controls and more peptidase activity, but less growth in bacteria exposed to CtenRNAV lysate compared to CtenDNAV lysate^20^, introducing the caveat that stimulation may not translate to growth or secondary production by the bacterial community.

#### Exploring the bioactivity of NVO vs PUAs

As mentioned above, NVOs are thought to be more deleterious to grazers than PUAs. For example, dinoflagellate grazing on diatom cultures is inhibited by 50% at 1 nM NVO but exposure to 10 nM PUA only decreased grazing by 33%; PUA concentrations in the 1000 nM range were needed for a 50% reduction in grazing^33^. To better understand the response to NVOs vs. PUA across multiple trophic levels, a set of grazing-growth experiments (Fig. 6I), two sets of surface microbial enzymatic activity experiments (Fig. 6J), and one set of particle-associated microbial enzymatic activity experiments (Fig. 6K) were conducted with the NVO, 15-HpETE, replicating the aforementioned PUA experiment. The grazing experiment with 15-HpETE (Fig. 6I) displayed large variability amongst replicates and a negative growth rate in the control and was inconclusive (Fig. S4D). Enzymatic activity was stimulated by NVOs in both surface and sinking particle-associated microbial communities (Fig. 6J-K). The stimulation of the surface community was more significant for NVOs than PUAs at 1 µM (Fig. 6J *vs.* 6E; St. 7). Similarly, particle-associated community responses to 10 μM PUAs and NVOs additions were observed at the Post-lytic Point Reyes sites (St. 8 and St.10, respectively; Fig. 6H and 6K). We had a limited amount of 15-HpETE stock, so we were restricted to a smaller number of experiments and a limited range of concentrations tested. Additional parallel experiments with NVOs and PUAs are needed to fully understand dose-dependent impacts of oxylipins on the same population. Nonetheless, our results are congruent with studies exploring the effect of various oxylipins on copepods, sea urchins, and dinoflagellate grazing, which suggest that the effective concentration varies by oxylipin^8,10,12,33,65,66^.

Notably, 15-HpETE, the compound tested in deck-board incubations, was observed in the DYEatom cruise meta-lipidome (Fig. S6A). HpETE analogs (*i.e.* compounds annotated as NVO 20:4 +2O at all retention times) were present in significantly higher concentrations at the Post-lytic Point Reyes sites compared to mid infection or non-bloom states (Fig. S6B). HpETE analogs represented less than 1% of the average annotated peak area (Fig. S7). C20:5 and C16:n oxylipins and their free fatty acid precursors dominated the dissolved lipidome representing 58% of the peak area (Fig. S6). Given the dose dependence of oxylipin bioactivity, oxylipin quantification is critical to understanding and contextualizing their bioactivity *in situ*. Our general untargeted approach better captured the wide diversity of compounds present but it only allows for presentation of relative abundances. Future work should take advantage of tractable absolute quantification with an adequate suite of representative authentic standards.

### Biogeochemical impacts of oxylipin chemical signaling during viral infection

We previously reported enhanced ectoprotease activity at the lytic Point Reyes site, which we interpreted as stimulation of remineralization and the ‘viral shunt’^17^. Deck-board experiments in this study revealed oxylipin-inducible enzyme activity in surface and sinking particle-associated microbial communities further connecting virus infection (and the induced chemical signaling) with enhanced organic matter turnover. Silicon dissolution is critically controlled by proteolytic degradation of the organic matrix protecting diatom frustules, denuding and exposing the opal (SiO_2_) to the undersaturated seawater^68^. In turn, this dissolved silicic acid (Si(OH)_4_) can support regenerative biogenic silica production^69^. Enhanced silica dissolution can also negatively impact mineral ballasting of exported carbon and diminish the oceanic silica pump^70^. We conducted experiments to test whether oxylipin enhanced enzymatic activity impacted biogenic silica dissolution, but it was inconclusive (data not presented). The stimulation of ectopeptidase in surface microbial communities has been observed in diatom blooms in this region^70^. The mechanisms underlying peptidase stimulation during viral infection of *Chaetoceros*^20^ are not well understood but could be linked to the production of nitrogen-rich, Coomassie Stainable Particles (CSP), which have been observed in virus-infected *Chaetoceros* culture studies^71^. Stimulation of peptidase over phosphatase in the surface ocean would serve to preferentially remineralize nitrogen from dissolved organic matter and is consistent with nitrogen being the co-limiting nutrient in the sunlit CCE along with Fe rather than phosphorus^72^. A preference for phosphatase stimulation on particles reflects a shift in community activity towards accessing organically bound phosphate in response to oxylipin chemical signaling below the euphotic zone.

At the same time, our geochemical data (Fig. 4A-C) strongly suggest that virus infection of this natural diatom community induced the viral shuttle and enhanced carbon export even with enhanced enzymatic turnover of organic material in surface communites^17^. Export was positively correlated with the total peak intensity of oxylipins in the dissolved lipidome (Fig. S5A), suggesting that oxylipins may play a role in enhanced export during infection. The relative balance between shunt versus shuttle outcomes will ultimately be determined by the remineralization length scales relative to the mixed layer depth (see Figure 5 in Kranzler et al.^20^). In our study, virus infection stimulated export flux and high e-ratios to 50 m; the observed MLD at this station was 10 m, thus driving the net outcome towards the shuttle.

Evidence of a strong viral shuttle was previously observed in coccolithophores populations that were actively infected with giant, double-stranded DNA-containing Coccolithoviruses in the North Atlantic^59^; diagnostic lipid biomarkers^48,73,74^ showed that sinking aggregates were enriched with infected cells and resulted in very high POC and PIC fluxes. Similarly, eukaryotic viral marker genes in a meta’omic study of the TARA Oceans Expedition global dataset explained 67% of the variability in carbon export efficiency with 58 viruses positively correlated and 25 negatively correlated with export efficiency^19^. Many of the positively correlated viruses were linked to silicifying eukaryotes, like diatoms. Mechanistically, culture studies with eukaryotic phytoplankton, including diatoms and coccolithophores, have shown that virus infection enhances spore formation^75^, TEP and CSP excretion, and particle aggregation ^71,76^. Oxylipins are known to increase aggregation in the benthic diatom *Fistulifera saprophila*^77^ but these effects are strain dependent.

In addition to remineralization length scales of sinking particles, population dynamics have been proposed as important controls on whether viral infection leads to enhanced export or upper ocean recycling. Large scale mesocosm experiments show a 2-4x increase in TEP and PIC production per cell for *G. huxleyi* during infection feeding into the viral shuttle and resulting in adsorption of viruses from the water column on to aggregated particles. Taken with the data presented here and the conceptualization of larger phytoplankton being associated with more zooplankton degradation and higher overall fluxes, we hypothesize that eukaryotic algae are more subject to the viral shuttle compared to prokaryotic hosts. The remineralization length scale plays into this as eukaryotic phytoplankton sink faster than prokaryotes as a function of size and added density due to associated biominerals; thus, the length scale over which their biomass can be remineralized is longer leading to more efficient carbon export. Our analysis adds a layer of complexity to the dynamic controls on viral carbon cycling with oxylipin-producing diatom populations altering organic matter recycling, grazing, growth, and export. The dose dependent nature of oxylipin signaling may contribute to episodic export once inhibitory thresholds for the community are surpassed, resulting in a toggle between shuttle and shunt (Fig. 7).

**Figure 7.**
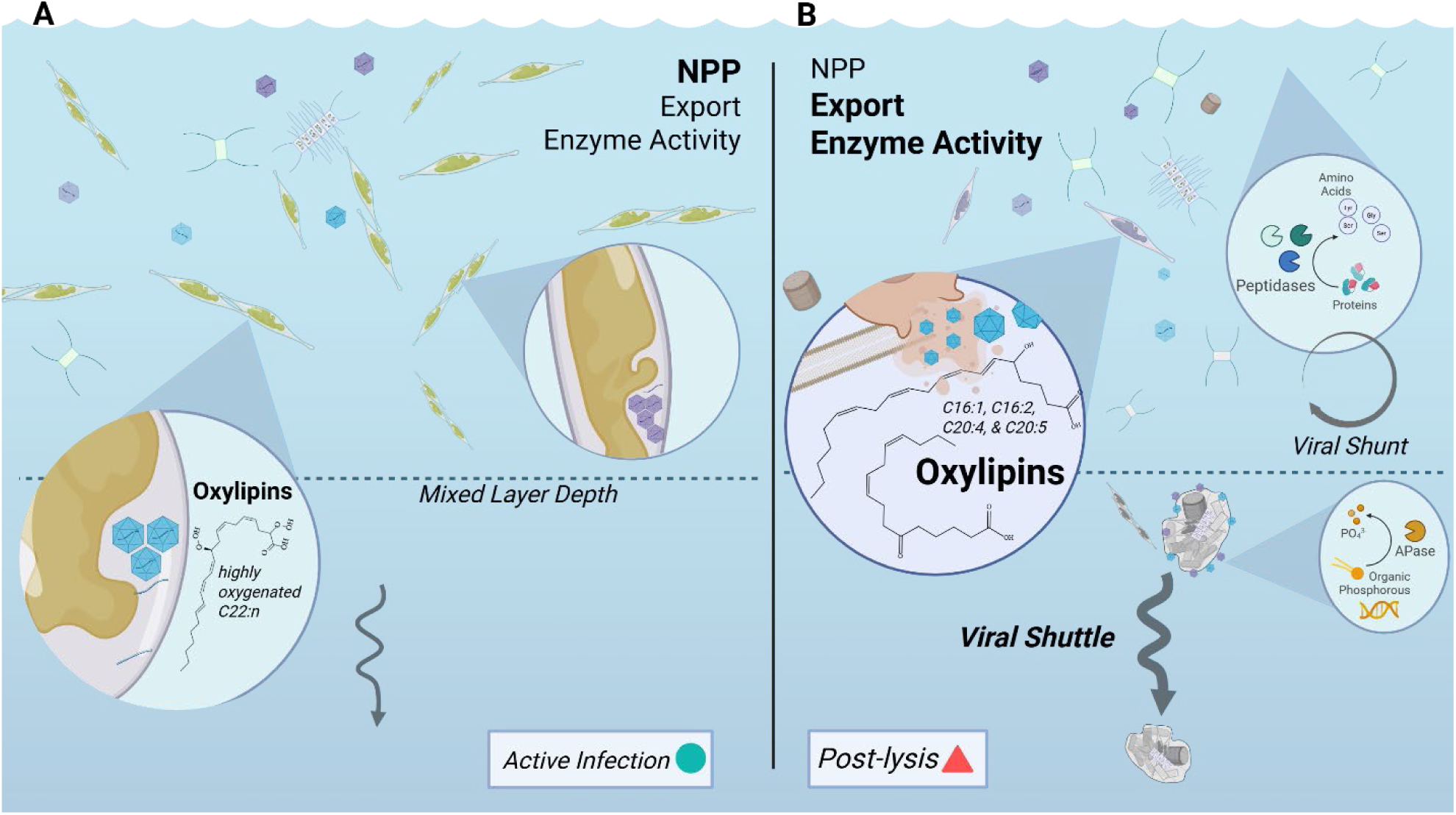
Oxylipin chemical signaling during virus infection and bloom demise toggles the fate of organic matter between export and remineralization. Conceptual link between the state of virus infection, oxylipin production, export flux, and remineralization length scale in a natural diatom population. **A)** Communities undergoing active infection (such as those surveyed near Monterey Bay on June 28^th^) were at the peak bloom NPP, as per satellite measurements and *in situ* chlorophyll accumulation. Some oxylipins are detected in the environment but at lower levels and were characterized by a different suite of compounds. **B)** Lytic populations (such as those surveyed at Point Reyes) had higher dissolved oxylipin concentrations, including those specifically viral biomarkers from culture infection studies, along with decreases in NPP over the proceeding 8 d, higher export flux and e-ratios of carbon to 50 m (compared to Monterey Bay), and higher ectoenzymatic activities. Taken together, these data suggest that virus infection and oxylipins induced the viral shuttle and enhanced carbon export even with enhanced turnover of organic matter remineralization indicative of the viral shunt^17^. The dashed horizontal line in each panel represents the mixed layer depth.

## Conclusions

Several, specific oxylipins were identified as biomarkers for viral infection and cell mortality in cultured *Chaetoceros* sp. and were subsequently found in the meta-lipidomes of virus-infected natural diatom communities infected in the California Current Ecosystem. Deck-board incubations agreed with microbial responses to oxylipin observed in previous studies, namely dose-, compound-, and community-dependent inhibition of grazing and phytoplankton growth and stimulation of particle-associated microbial communities. Oxylipins stimulated phosphatase activity on sinking particles and peptidase activity in surface communities, while inhibiting microzooplankton grazing and phytoplankton growth. Biogeochemical measurements suggest that virus-mediated diatom bloom decline led to higher and more efficient carbon export compared to earlier phases of infection, despite stimulatory effects on enzymatic remineralization of organic matter. Taken together, these findings highlight how viruses and chemical signaling via oxylipins mediate trophic interactions and biogeochemical fate of organic matter in productive upwelling regimes.

## Materials and Methods

### Viral Infection Experiments with Cultured Chaetoceros sp

Cultures of the diatom hosts *C. socialis* strain NIES-3713 (formerly L-4) and *C. tenuissimus* strains NIES-3714 (formerly 2-6) and NIES-3715 (formerly 2-10) were grown in SWM-3 media at 15°C under 150 µmols photons m^-2^ s^-1^ irradiance on a 12:12 L:D photoperiod. Triplicate cultures were infected during exponential growth with CsfrRNAV (*C. socialis*), CtenRNAV (*C. tenuissimus* NIES-3715) or CtenDNAV (*C. tenuissimus* NIES-3715). Samples were collected for cell counts via microscopy or flow cytometry. For experiments with CsfrRNAV and CtenRNAV, samples for dissolved lipidome analysis were collected at 0, 1, and 3 dpi, representing early, mid, and late infection, respectively (Fig. S2). For experiments with CtenDNAV, samples for dissolved lipidome analysis were collected 0, 1, 3, and 4 dpi. Triplicate uninfected control cultures were sampled concurrently to exclude changes during normal growth. Follow-up experiments grew the two hosts, along with *C. tenuissimus* NIES-3714 into decline phase to assess changes associated with death (likely induced by nutrient limitation). Samples were pre-filtered through a 0.2 μM Durapore filter to remove particles. The deuterated internal standard, 15(S)-Hydroxy eicosatetraenoic acid-d8 (Cayman Chemical), was added to each filtered sample to a final concentration of 10 μM. The dissolved lipids were then extracted onto a Waters HLB SPE cartridge.

#### Lipidomic Analysis

The dissolved lipids were eluted from the SPE cartridge with 2 ml of 70:30 acetonitrile: isopropanol into pre-combusted collection vials that were treated with the antioxidant BHT to prevent autoxidation of the samples. The samples were immediately transferred from the collection vials into HPLC vials and capped under argon. The samples were stored at -80°C until mass spectrometric analysis. Within a week of extraction the dissolved lipidome samples were analyzed using reverse phase HPLC (Agilent 1200 system; Agilent, Santa Clara, CA, USA) paired with high resolution, accurate mass (HRAM) data from a Thermo Exactive Plus Orbitrap mass spectrometer (ThermoFisher Scientific, Waltham, MA, USA). The chromatographic method used a Xbridge C8 column (Waters, Milford, MA, USA) as the stationary phase, 18 MΩ water as eluent A, 70:30 acetonitrile: isopropanol as eluent B, and ammonium acetate and acetic acid as the adduct forming additives^70^. The gradient began with a 1-minute isocratic hold of 45% B and shifted to 99% B over 25 minutes with a flow rate of 0.4 mL min^-1^. Full scan data (m/z 100-1500) was collected in negative mode at a mass resolution of 170,000. For the follow-up experiments and to annotate the oxylipins in the original Chaetoceros and DYEatom cruise samples, the dissolved lipid extracts were analyzing using reverse phase chromatography (Vanquish UHPLC, ThermoFisher Scientific, San Jose, CA, USA) paired with HRAM data-dependent acquisition on a Orbitrap ID-X mass spectrometry (ThermoFisher Scientific, San Jose, CA, USA). A mass inclusion list with the exact masses of [M-H]^-^ adducts of important oxylipins was used to acquire ms^2^ for further annotation (*Chaetoceros* experiment oxylipin annotation described in Edwards et al. 2024; DYEatom annotation in Table S5). For further details, please refer to the manuscript detailing the intercomparison of diatom host-virus pairs^16^.

#### Chemoinformatic Pipeline

The raw mass spectrometric data was converted to mzXML format with mscovert and analyzed with the R packages xcms^78–80^ and CAMERA^81^, which aligned the chromatograms, picked and integrated peaks, and identified and removed secondary isotopic peaks (specific details in Edwards *et al*., 2024). The lipidomic features were further annotated using the LOBSTAHS package which included m/z information of the various acyl chain-lengths, degrees of unsaturation, and oxidations of triacylglycerols, free fatty acids (FFA), polyunsaturated aldehydes (PUA), and eight intact polar diacylglycerol lipids and their most common adducts^82^. The features annotated as oxylipins, and free fatty acids were retained and manually verified as high-quality peaks in El-MAVEN^83–85^. The fragmentation of the compounds was assessed using the rerun of the 2016 samples on the UPLC-Orbitrap ID-X and in silico databases created by the Edwards Lab in CFM-ID^86^, and iterative database of mammalian oxylipins from Watrouse^87^, and the *in silico* database of algae, plant, and animal lipids available through the LipidBlast pathway in MS-DIAL^88^. Peak areas were normalized to the recovery of the internal standard and volume filtered. Statistical analyses were performed using the R package MetaboAnalyst through its web interface^54^.

### DYEatom cruise

The DYEatom cruise aboard the R/V Point Sur (PS1312) disembarked from Moss Landing, CA on June 27, 2013. The 9-day cruise track transected the California coastal upwelling region (Fig 4 and Table S4). Nine stations were occupied. The details of the metatranscriptomic analysis of cell associated viruses and qPCR of free-living viruses used for assessing viral infection by diatom RNA viruses can be found in Kranzler *et al.* 2019, where the details of the enzyme activity protocols can also be found^17^.

#### Dissolved Meta-lipidome

The dissolved lipidome was sampled at two depths in the water column along the cruise track, the 55% Io depth and the chlorophyll maximum. One liter of seawater from each depth was filtered through a bottle top 0.2 μm filter (Corning) to remove particulate material. The internal standard benzaldehyde (Sigma-Aldrich) was added to a final concentration of 10 μM in one liter of seawater. Using a vacuum manifold, the samples were filtered onto a Water HLB solid phase extraction (SPE) cartridge for approximately one hour and the final volume filtered was recorded. The SPE cartridges were acidified with 0.1% hydrochloric acid, placed in a whirl-pak bag, and stored at -80°C until elution and analysis at Woods Hole Oceanographic Institution. Samples were analyzed using the chromatographic, mass spec, and lipidomic pipeline outlined above.

#### Sediment Trap Collection of Sinking Particles

At Stations 1, 6, 8, and 10, sinking particles were collected using a surface tethered net trap deployed at a depth of 50m for ∼8 hrs. Trap material was screened through a 350 µm mesh to exclude larger mesozooplankton and split into 8 equal 500 mL fractions using a rotating electric splitter^89,90^. One 500-mL split of the trap material was filtered onto three pre-combusted 47mm glass fiber filters (nominal pore size-0.7-μm) for particulate organic carbon and nitrogen (POC/PON) analysis. The samples were kept at −80 °C until analysis at Woods Hole Oceanographic Institution (Woods Hole, MA) on a Finnigan-MAT DeltaPlus stable light isotope ratio mass spectrometer^91^. The POC and PON values of the three filters were summed together for each trap and deployment time was used to calculate flux to 50 m over a 24-hour period.

#### Net Primary Productivity Estimates

High-resolution, 8-day integrated satellite-based estimates of net primary productivity in the CCE for the month surrounding our cruise were downloaded from the Oregon State University Ocean Productivity website. The NPP estimates come from the CAFE model, which is depth, time, and spectrally resolved^55^. Clara Douglas wrote the MatLab script for this analysis.

### Deck-board Incubations

Three types of amendment experiments were conducted at sea: 1) three deck-board incubation experiments with surface seawater communities and varying concentrations, of either a mix of PUAs (0, 0.1, and 1μM) or the NVO, 15-HpETE (0, 0.1, and 1 μM; Fig. 6B), 2) four dark incubation experiment with sinking particles from 50 m amended with 0, 1, and 10 μM HpETE or 0, 1, 10, and 100 μM PUA (Fig. 6C); 3) seven deck-board paired dilution experiments to compare microzooplankton grazing rates and phytoplankton growth rate in no amendment controls and 1 μM PUA or 0.8 μM HpETE treatments (Fig. 6A). These concentrations were chosen based on the bioactive PUA concentrations for marine bacterial isolates^28^, PUA concentrations that enhanced the metabolism of marine microbes associated with sinking particles in the North Atlantic^35^, and PUA concentrations that decreased microzooplankton growth rate in the Chesapeake Bay^32^.

#### Stock solutions

The NVO 15-HpETE stock solution was biosynthesized in the lab by adding the enzyme 15-lipoxygenase (Sigma-Aldrich) to arachidonic acid (Sigma-Aldrich) using the protocol outlined in Iacazio *et al.* (1990)^92^. Once the reaction reached completion, the oxylipins were extracted onto an SPE cartridge (HBL Waters) and immediately eluted with methanol. Mass spectrometric analysis determined the composition of the oxylipin stock was 825 mmol L^-1^ 15-HpETE and a background of its precursor and positional isomers.

The oxylipin stock solution was capped under N_2_ and stored at -20 °C. In our LOBSTAHs notation HpETE is FFA 20:4 +2O.

A stock solution containing a mix of polyunsaturated aldehydes was prepared at sea and contained 100 mM of 2,4-heptdienal, 2.4-octadienal, and 2,4-decadienal (Sigma-Aldrich) dissolved in methanol. The solution was kept at 4C and was remade every two days to prevent degradation. In our LOBSTAHS notation these compounds would be PUA 7:2, PUA 8:2 and PUA 10:2.

#### Surface Ocean Community

At Stations 4, 7, and 10, seawater was collected from the 55% I_0_ depth into 2 L Nalgene bottles. At St. 4, 0, 0.1, 1 and 10 μM PUAs were added to triplicate bottles using methanol as a carrier liquid. Care was taken such that the same volume of methanol was added to each treatment including the 0 μM control. At St. 7, 0, 0.1, and1 μM PUA and 1 μM HpETE were added to bottles in triplicate. At St. 10, we began to run out of oxylipin stocks so only three conditions were made up in triplicate: 0, 0.1, and 1 μM HpETE. The 2 L microcosms were placed in on-deck incubators screened to mimic the light field at 55% I_0_ and temperature controlled by a flow through system for ∼24 hours. Enzymatic activity was measured at the end of the experiment (Fig. 6B) using fluorogenic substrates as described in Kranzler *et al.* 2019^17^.

#### Particle Associated Community

At the stations where sinking particles were collected, a 500 mL split was used to set up dark incubations using the experimental design laid out in Edwards et al 2015. In a dark on-deck incubator that was temperature controlled by a flow-through system, the trap material was incubated in opaque 150 mL Nalgene bottles in triplicate for 24 hours. At St 1, 6, and 8, the treatments were 0, 1, 10, and 100 μM PUA. At St. 10, the treatment was 0, 1, or 10 μM HpETE Enzymatic activity was measured at the end of each experiment (Fig. 6C). Samples were also taken for biogenic silica quantification by filtering 25 mL onto a 1.2 μm nucleopore filter for analysis at Rutgers (Fig. S8A). The same volume was filtered through a 0.7 μm nucleopore and frozen for dissolved Si analysis at UCSB (Fig. S8B).

#### Grazing and Growth Rate-Dilution Experiments

Paired dilution experiments were conducted to measure microzooplankton grazing rates and phytoplankton growth rates with and without exposure to exogenous oxylipins, at seven stations along the cruise track (Fig. 6A and S4). At each station, 40 L of water was collected from the depth of 50% surface photosynthetically active radiation (PAR). Half of this volume was then gravity filtered through a 200 μm Nitex mesh, to be used as whole seawater (WSW), and stored in an on-deck incubator at *in situ* surface temperature until the experimental set-up. The other 20 L was gravity filtered through a 20 μm Nitex mesh, to gently remove large phytoplankton (i.e. diatoms), and then gravity filtered through a Pall 0.2 μm vented sterile filter capsule, in order to make dilution water. Once the dilution water was prepared, we used a modified dilution method^93^ with three points, using 5, 20, and 100% WSW, in triplicate 1L bottles. Nutrients were not added to bottles. For treated bottles, a final concentration of 0.8 μM oxylipins were added using a stock solution prepared in the lab. All dilution bottles were sampled at time 0 and after 20-24h for chlorophyll, cell counts (1% gluteraldehyde), and FIRe analysis, while only 100% WSW bottles were sampled at both time points for microzooplankton abundance (10% Lugol’s).

#### Statistical analyses for incubation experiments

For the oxylipin amendment experiments, Shapiro-Wilk tests showed that the data was not normally distributed and non-parametric rank-order tests were used to determine significant differences between treatments (Sigma-Plot). A Mann-Whitney U was utilized to test whether microzooplankton grazing rate and the phytoplankton growth rates from the paired dilution experiments were significantly different in the no amendment controls compared to the 0.8 μM HpETE treatments. A Kruskal-Wallis analysis of variance on ranks was used to determine if the enzymatic activity of peptidase and phosphatase was significantly different in the controls compared to the oxylipin treatments in amendment experiments, as well as, to determine if the total peak area of annotated free fatty acids and oxylipins was significantly different between the significant groups of samples revealed by similarity profile analysis (p < 0.1).

Dunn’s method was used as a post-hoc to determine which groups were different from one another (p < 0.05).

## Supporting information

Supplemental File 1. Chaetoceros viral infection dissolved lipidome

Supplemental File 2. DYEatom cruise dissolved meta-lipidome

Supplemental File 3. Retention time model for DYEatom lipidome annotation

## Data Accessibility

The dissolved lipidomes are available on the MassIVE repository. Biogeochemical data from the DYEatom cruise can be found on BCO-DMO (PS1312, http://www.bco-dmo.org/project/550825).

## Acknowledgements

The captain and crew of the *R/V Pt. Sur* and the Brzezinski lab group who also sailed with us. We thank Yuji Tomaru (Fisheries Technology Institute, Japan Fisheries Research and Education Agency) for providing the *Chaetoceros* diatom host-virus systems. We thank the funding sources that supported this work: NSF OCE-1155663 (to JWK), Gordon and Betty Moore Foundation (BVM, KDB, MJ), McMinn Endowed Fund (BRE), NSF OCE-(BRE),

## Supplemental Information

**Figure S1.**
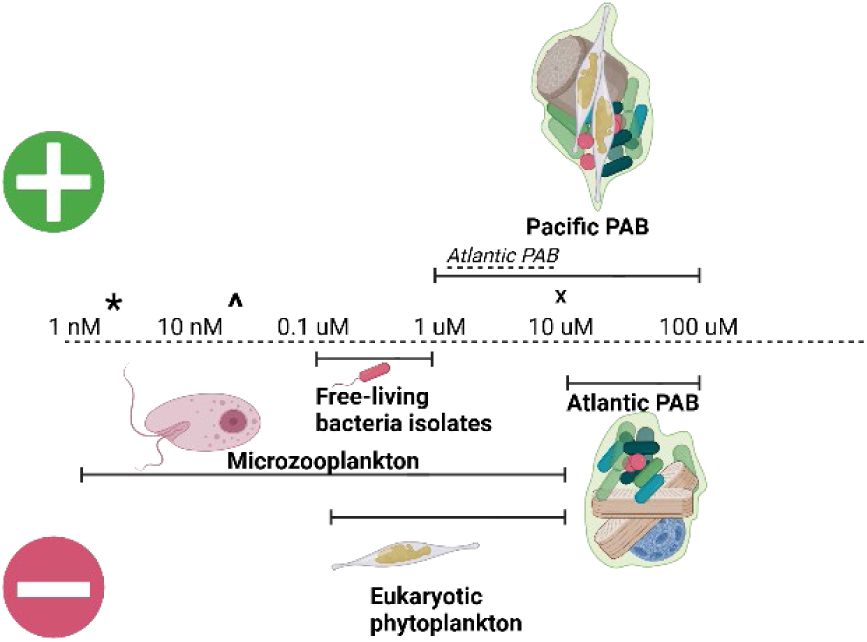
A summary of the range of concentrations known to be bioactive in amendment experiments and the maximum concentrations observed across the global ocean. The asterisk is the highest dissolved PUA concentration observed *in situ.*^94^. The caret is the highest particulate PUA concentration observed *in situ*^94^. The ‘x’ is the highest PUA concentration observed within sinking particles^35^.

**Figure S2.**
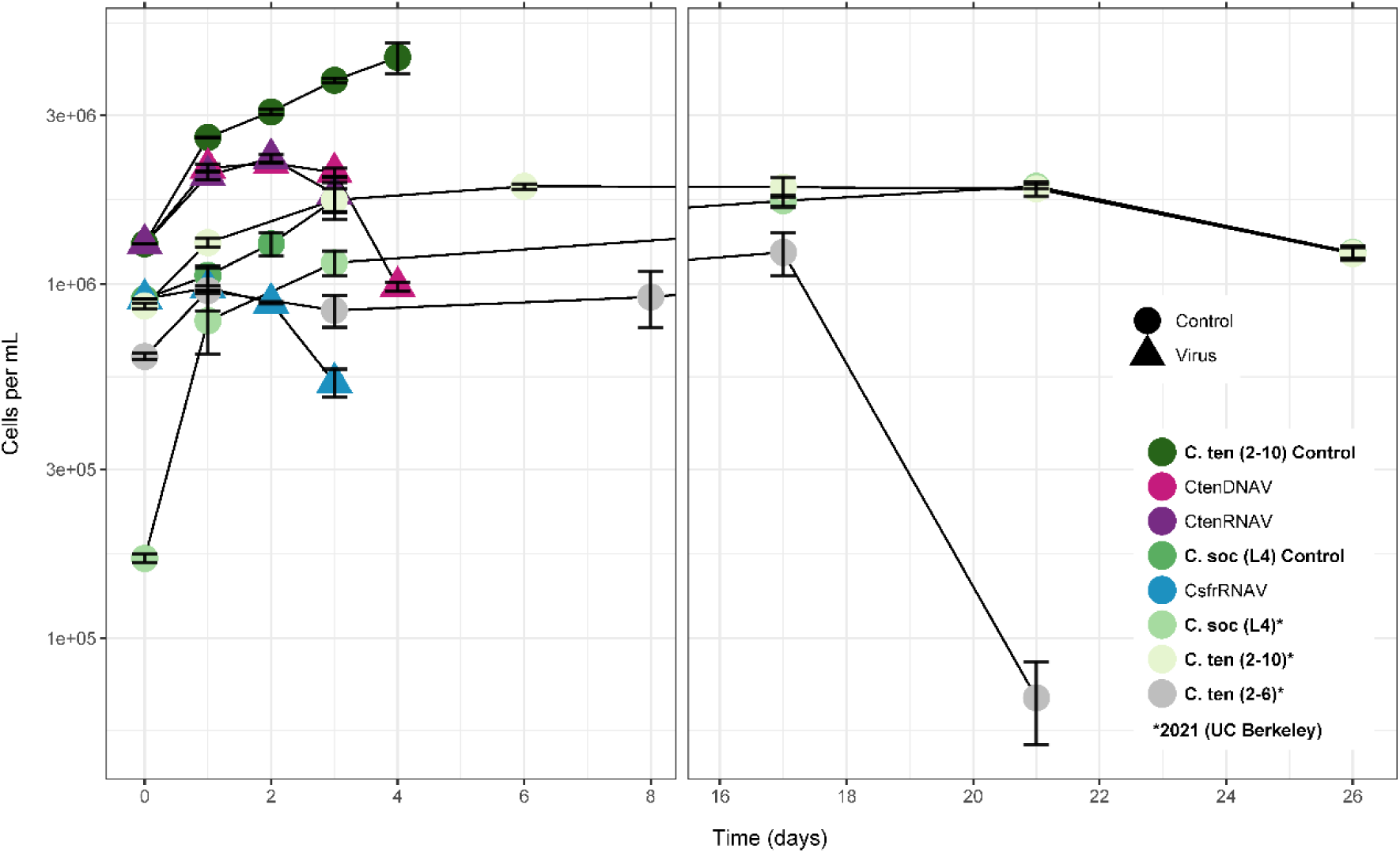
Time course of virus infection of *C. tenuissimus* and *C. socialis*. Growth curves for uninfected host-only control triplicates (green circles), *C. tenuissimus* infected with CtenRNAV (purple triangles) and CtenDNAV (pink triangle) viruses, and C. socialis infected with CsfrRNAV (blue triangles). Initial experiments at Rutgers in 2016 followed the diatom hosts, *Chaetoceros socialis* L4 (grass green circles) and *Chaetoceros tenuissimus* 2-10 (darkest green circle), to late exponential/ stationary phase and the dissolved lipid extracts were processed at WHOI. Subsequent experiments at UC-Berkely in 2021 followed cells well into stationary and decline phase (mint green, light green, and grey circles, denoted by an asterisk in the legend) were analyzed at UC-Berkeley.

**Figure S3.**
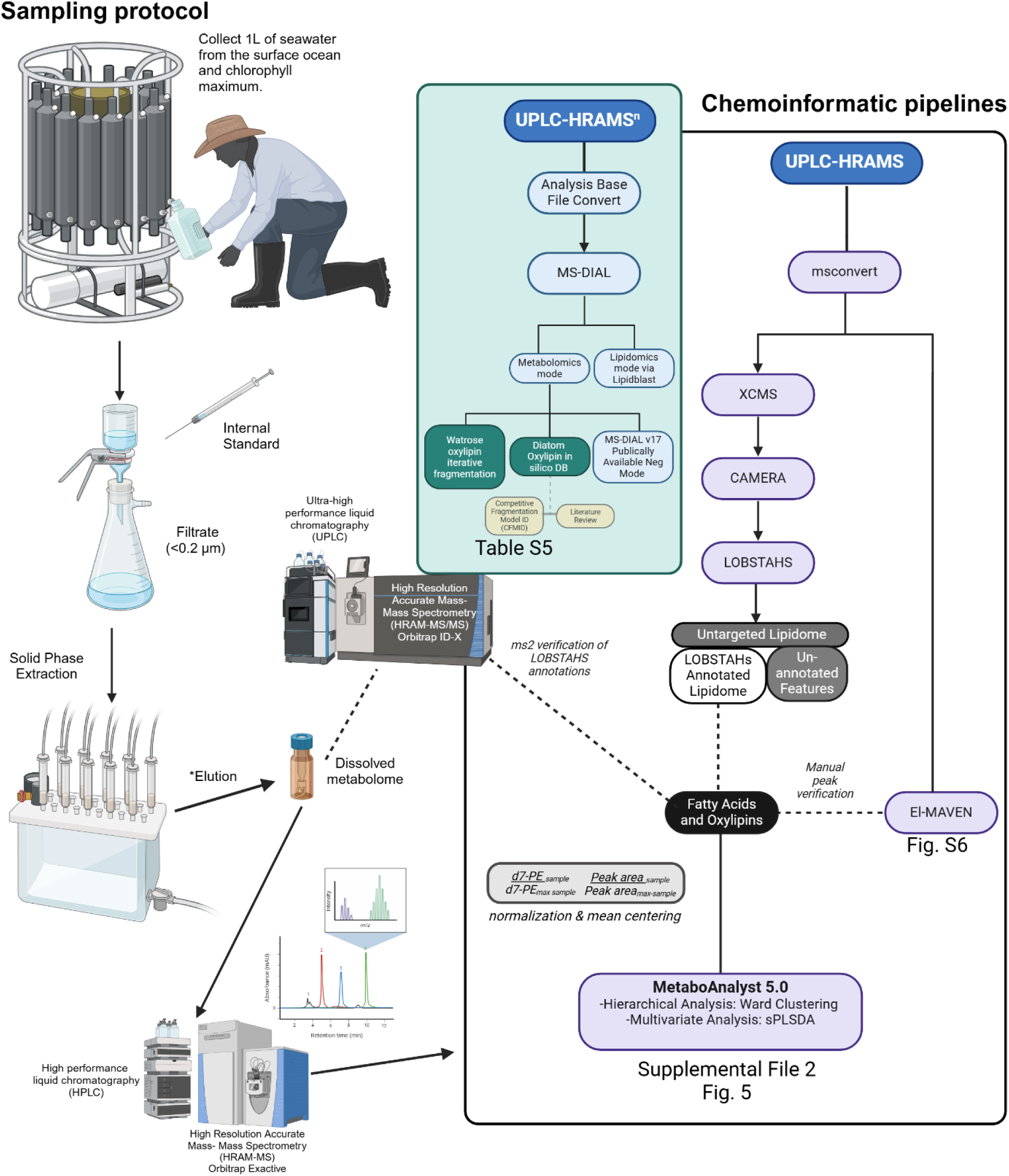
Outline of dissolved metabolome sampling at sea as well as sample extraction and analysis in the lab. One liter of water was collected from the surface ocean (55% I0 depth approx. 5 m) and the chlorophyll maximum (4-35 m) at each station. The 1L sample was filtered through a 0.2 um filter and benzaldehyde was added to the filtrate as an internal standard. The dissolved organics were extracted from the filtrate onto a Water HLB Solid Phase Extraction cartridge using a vacuum manifold. SPE Samples were flash frozen, and the dissolved metabolome was eluted from the SPE back in the lab. Dissolved metabolome samples were run for lipidomic analysis using HPLC paired with Orbitrap Exactive mass spectrometer in early 2014. Three samples were excluded from further analysis due to missing internal standard. The eleven remaining raw lipidomes were processed through the xcms-CAMERA-LOBSTAHS-MetaboAnalyst Chemoinformatic pipeline in the black box. Representative samples were rerun in 2021 using an Orbitrap ID-X mass spectrometer which allowed for more targeted ms2 fragmentation for higher confidence oxylipin annotation of important features from the comparative statistical analysis provided by MetaboAnalyst; these annotations can be found in Table S5.

**Fig S4.**
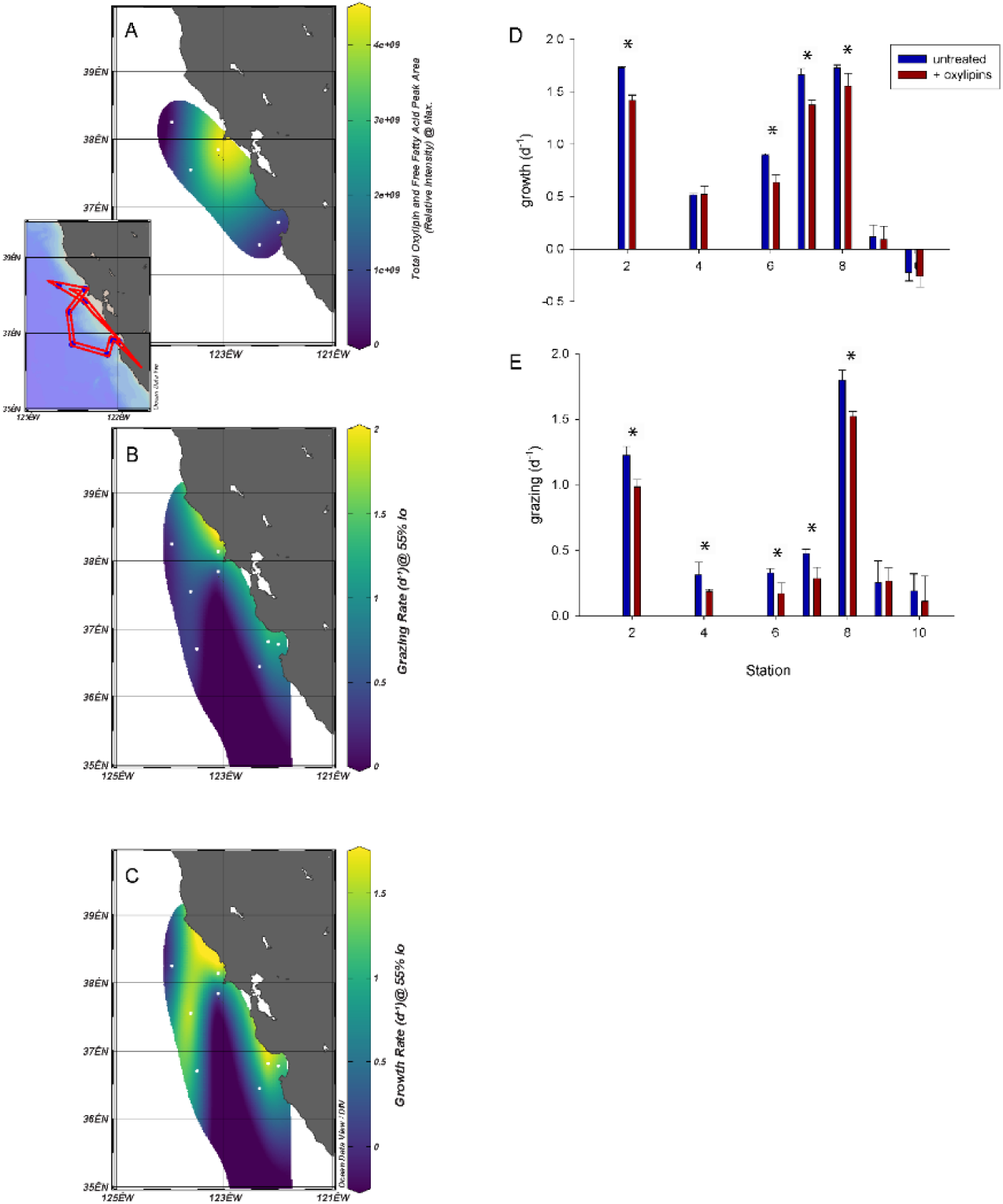
Grazing and growth were lowest at the station with the highest oxylipin concentration and deckboard amendment experiments with exogenous oxylipins show inhibition of growth and grazing. Contour maps of **A)** the maximum total oxylipin and free fatty acid peak area, **B)** microzooplankton grazing rate at the 55% I0 depth, and **C)** phytoplankton growth rate at the 55% I0 depth for each station of the DYEatom cruise. Plotted with Ocean Data View using the DIVA contouring model which accounts for general ocean current patterns. D) Grazing rate presented as a bar chart with the parallel oxylipin amendment rate. Asterisk denotes statistical difference in + oxylipin amendment from the control treatment. E) Growth rate presented as a bar chart with the parallel oxylipin amendment rate. Asterisks denote statistical difference from the control treatment.

**Fig S5.**
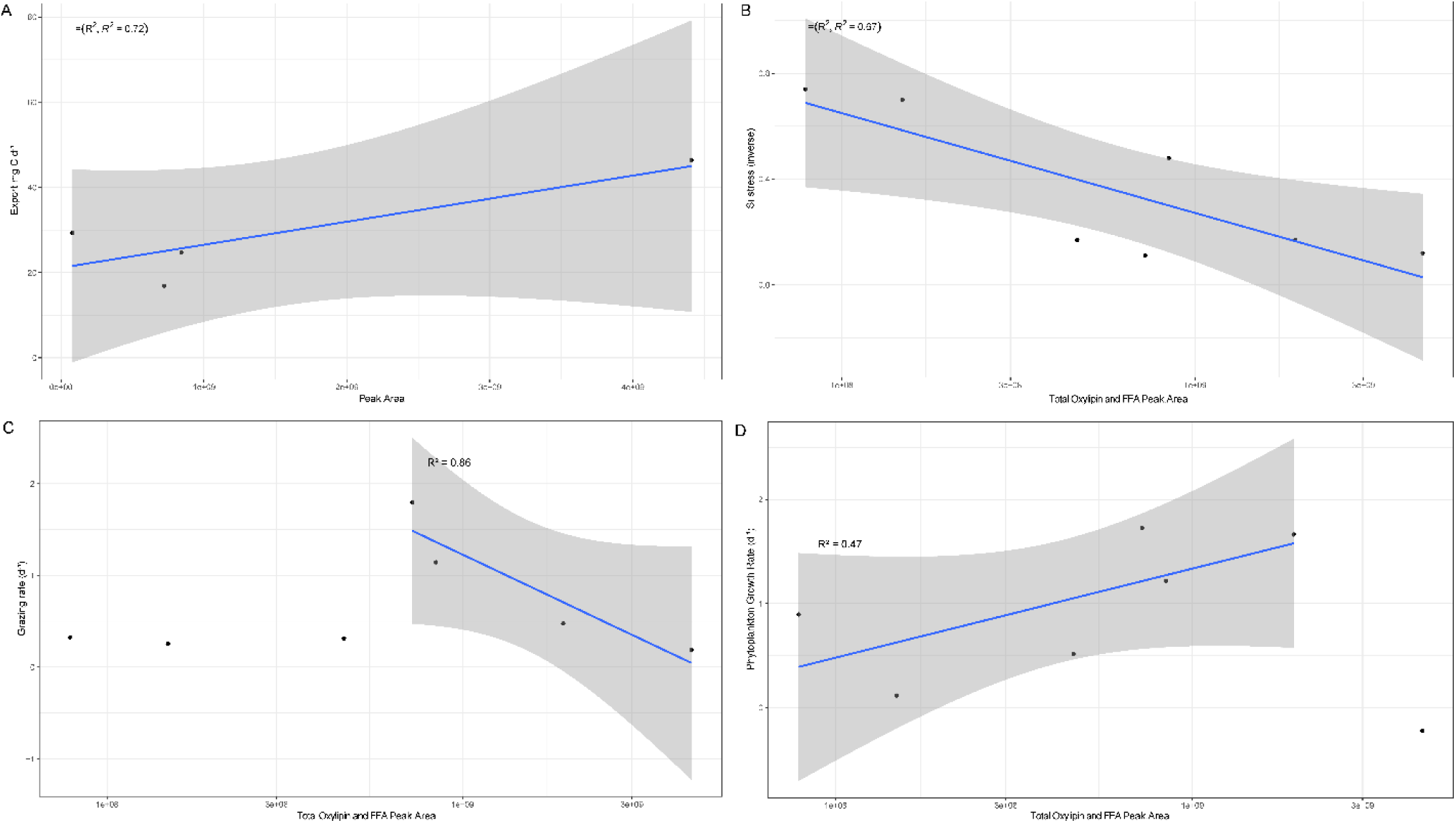
Relationship between the total peak area and **A)** export, **B)** Si-stress of the surface ocean community **C)** Grazing rate, and **D)** Phytoplankton growth rate across the DYEatom cruise (PS1312) reflect allelopathy of oxylipins on the community, oxylipin production under stress, and evidence of a link between oxylipin production and the viral shuttle. The silica stress metric is described in ref^17^. Briefly, the Y-axis is the percentage of the Si uptake rate (B) that happens in the ambient Si concentration vs. a deck-board amendment with exogenous Si. A lower value indicates more silicon stress in that environment. For the phytoplankton growth regression (D) all but the negative growth rate was included. And for the microzooplankton grazing rate regression (C) the low peak area samples were excluded.

**Figure S6.**
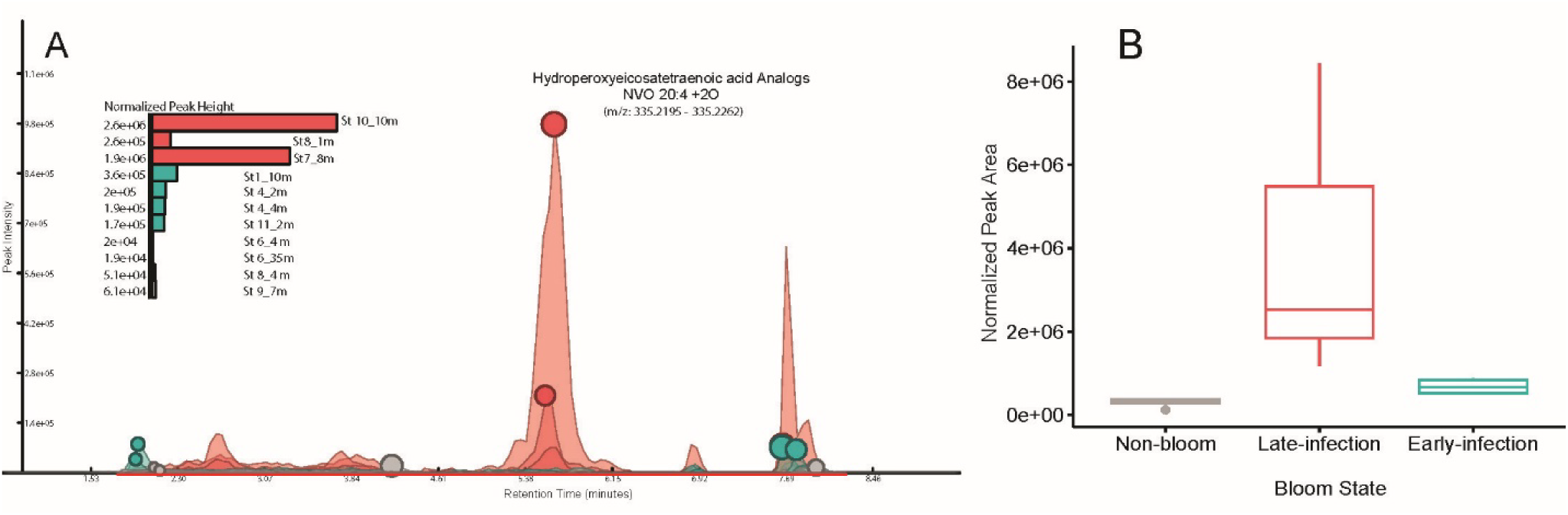
Analogs to the exogenous oxylipins found to be bioactive in Fig 6I-K were found *in situ* in addition to the various similar oxylipins of different carbon chain lengths and degrees of unsaturation highlighted in Fig 5. **A)** Extracted ion chromatogram of HpETE analogs (i.e. compounds with a [M-H]^-^ adduct m/z of 335.2232) observed in the DYEatom dissolved meta-lipidome and the integrated peak height displayed in the inset bar charts. **B)** The normalized peak area for HpETE analogs averaged across each bloom state. Box plots show the 1^st^ and 3^rd^ quartile range as the upper and lower bounds of the box and the mean is denoted as a line through the box. The error bars denote the standard deviation with the dots representing outliers.

**Figure S7.**
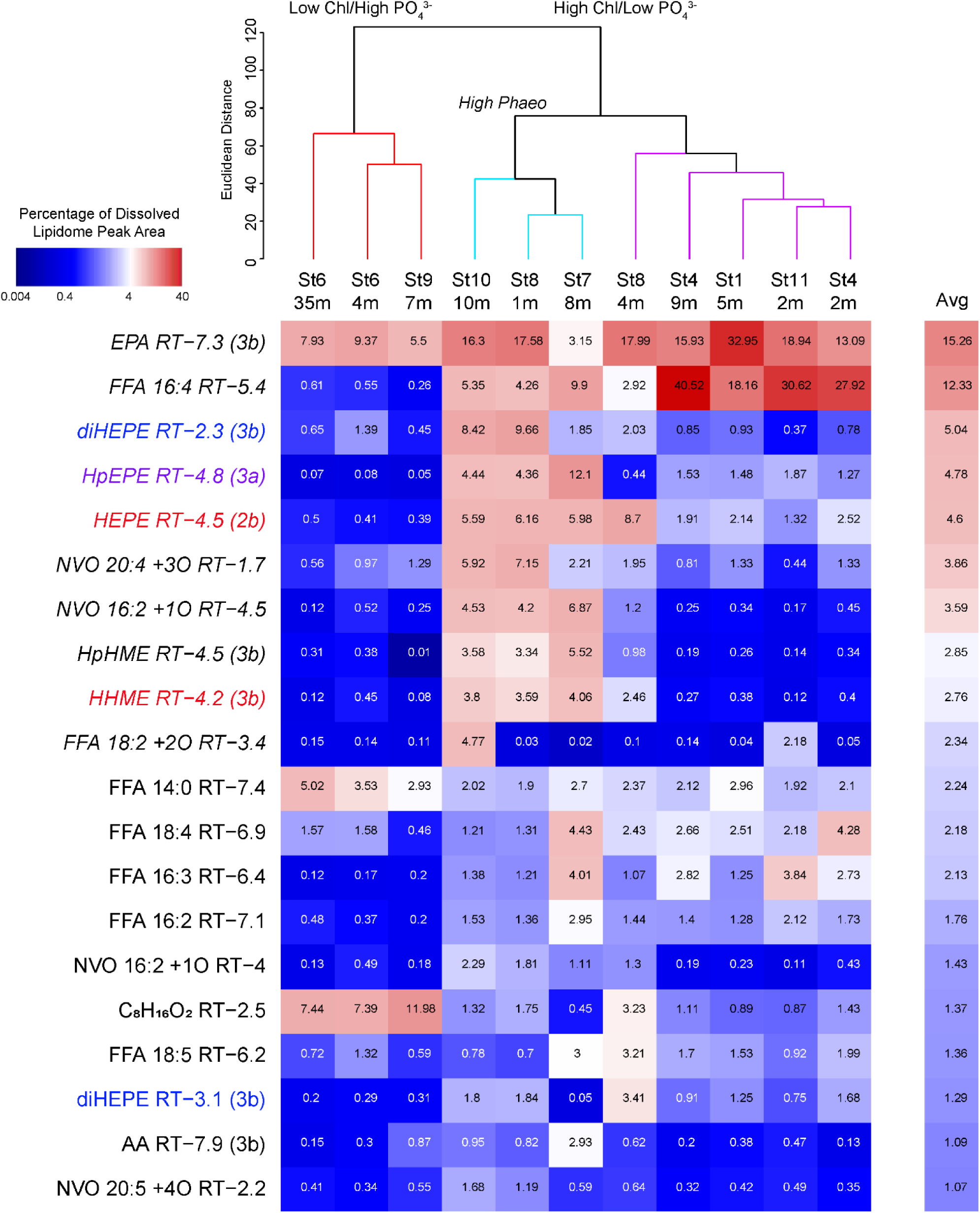
Features representing greater than 1% of the peak area in the annotated dissolved meta-lipidome on average. The heat map displays the percentage of the total annotated peak area attributable to each molecular species across the samples. Samples are grouped by similarity profile analysis of the structure of the dissolved meta-lipidome across the cruise track as presented in Fig. 5A.

**Table S1.**
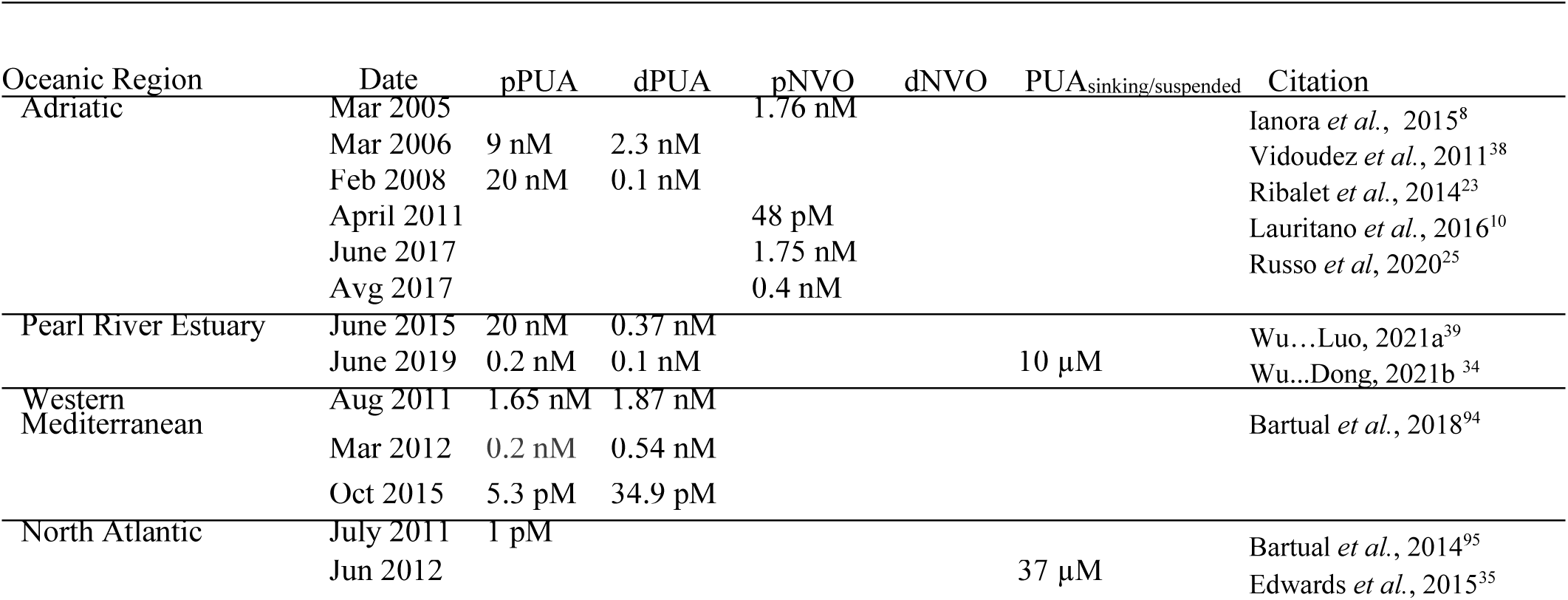
Compilation of maximum PUA and NVO concentrations observed in situ. Each oxylipin subclass is broken down into particulate (p-prefix) and dissolved (d-prefix). Note that no dNVO quantifications have been published. The pPUA values are derived from sonicated or freeze-thawed cells that were collected on a filter presenting a potential oxylipin concentration. The dPUA values were collected by extracting the organics on to a solid phase extraction cartridge. The pNVO values in Mar 2005 and April 2011 were estimated from cell abundances and fg of oxylipin per cell.

**Table S2.**
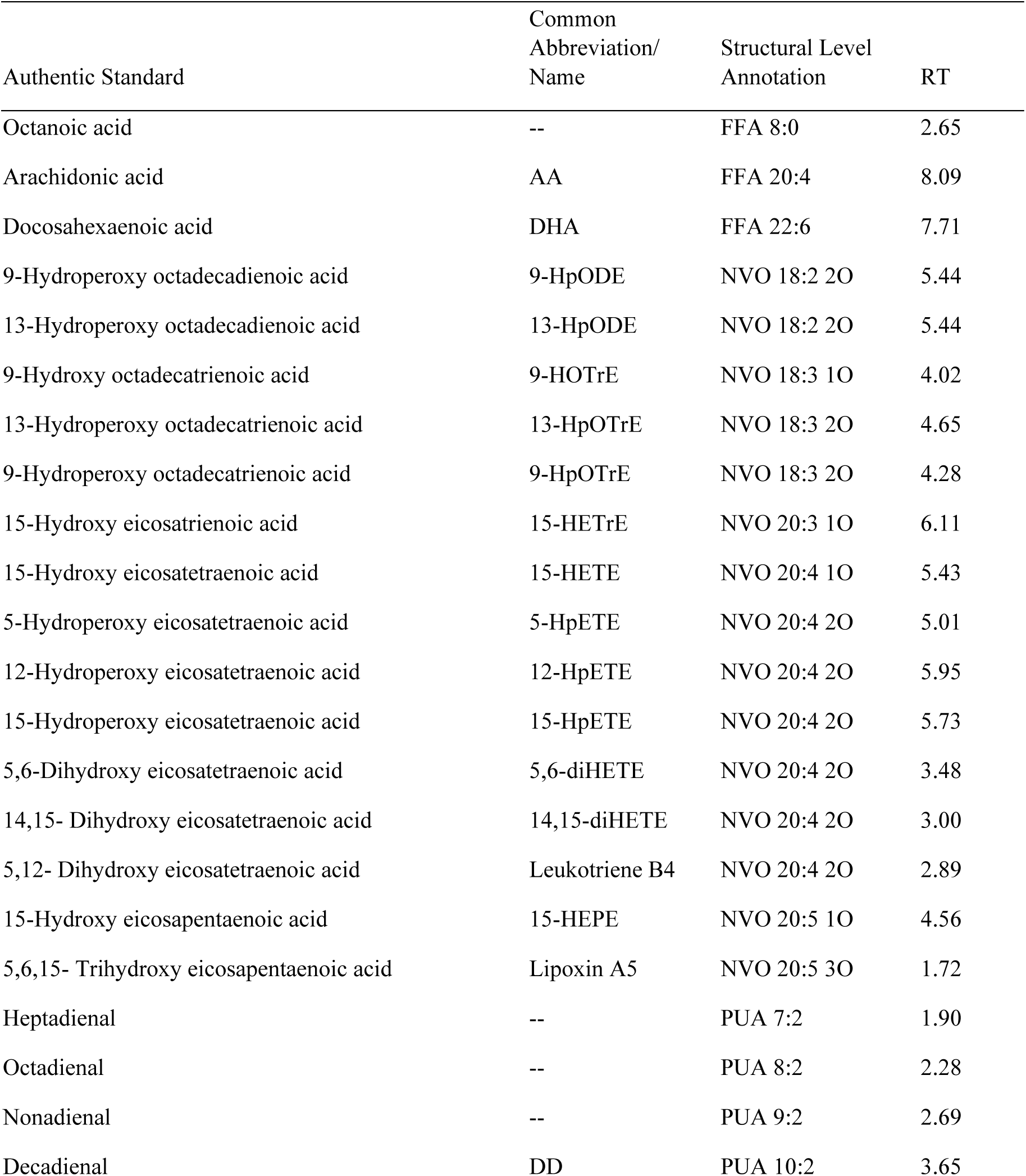
Authentic fatty acid, polyunsaturated aldehyde, non-volatile oxylipin standards with the common name or abbreviation, the structural level annotation assigned by the lipidomics pipeline _1022_ presented here, and the retention time (RT) of each compound. Comparisons to the RT of the authentic standards were used to assign putative annotations to features in the dissolved lipidome.

**Table S3.** Viral Infection and general death oxylipin-biomarkers from our *Chaetoceros*-virus experiments that were observed in culture, field studies, and their known allelopathy. The diatom species that each compound has been observed in is noted: *Skeletonema marinoi* (Sm), *Chaetoceros socialis* (Cs), *Chaetoceros affinis* (Ca)^96^, *Thalassiosira rotul*^97^ (Tr), *Pseudo-nitzchia delicatissima* (Pd)^98^, *Skeletonema pseudocostatum* (Sp)^13^, and *Phaeodactylum tricornutum* – microzooplankton grazing experiments (Pt-MZP)^33^. The lipidomic pipeline in this study automatically annotates to the structural level. Structural isomers of these compounds were observed in the DYEatom cruise dissolved lipidome (denoted as this study) and studies in the Mediterranean^8,10,25,99^. Compounds with known allelopathic effects are ^10,25^. Many of these compounds were annotated to the function group level using a retention time model or *in silico* fragmentation model as outlined in Edwards *et al.* (2024); compound abbreviations in Fig 3B-C

| Viral Biomarkers | Observed in culture | Observed in the Field | Known allelopathy | RT-model | CFMID model |
| --- | --- | --- | --- | --- | --- |
| FFA 14:2 +1O_mz_239.165_RT_1.71 | isomer Pt-MZP |  | MZP grazing |  |  |
| FFA 20:5 +2O_mz_333.2073_RT_2.08 | Sm, Ca, isomer Pt-MZP |  | Copepods; 20:4 +2O dihydro and hydroperoxy decrease MZP grazing; HpETE modulate particle associated microbial activity (this study) | 10,11-diHEPE |  |
| FFA 16:3 +3O_mz_297.1708_RT_2.13 | 16:3 oxylipins general Sm, Tr |  |  |  |  |
| FFA 14:2 +1O_mz_239.165_RT_2.56 | isomer Pt -MZP |  | MZP grazing |  |  |
| FFA 16:3_mz_249.1857_RT_7.36 | 16:3 oxylipins general Sm, Tr | Lauritano *et al.*, 2016 |  |  |  |
| FFA 22:5 +1O_mz_345.2436_RT_7.67 | *Lepidocylindrus* | Lauritano *et al.*, 2016; Russo *et al.*, 2020; this study | Copepods |  |  |
| FFA 14:2 +1O_mz_239.165_RT_2.25 | isomer Pt -MZP |  | MZP grazing |  |  |
| FFA 22:4 +1O_mz_347.2593_RT_7.77 | *Lepidocylindrus* |  |  |  |  |
| FFA 18:3 +2O_mz_309.2074_RT_5.01 | cyanobacteria, green algae, moss |  |  |  |  |
| FFA 20:4 +1O_mz_319.2279_RT_5.95 | Tr, *Stephanopyxis turris*, red algae, fungi | Lauritano *et al.* 2016; Russo *et al.* 2020; this study | Copepods; analog impacts sea urchins |  |  |
| Death Biomarkers | Observed in culture | Observed in the Field | Known allelopathy | RT-model | CFMID model |
| FFA 16:1 +1O_mz_269.2123_RT_4.37 | Sm, Tr | Lauritano *et al.* 2016; this study | Copepods |  |  |
| FFA 18:3 +2O_mz_309.2073_RT_1.98 | cyanobacteria, green algae, moss |  |  |  |  |
| FFA 16:2 +1O_mz_267.1966_RT_4.78 | Sm, Tr | this study | Copepods | 9-KHME | 9-epHME |
| FFA 16:3 +3O_mz_297.1707_RT_1.52 | 16:3 oxylipins general Sm, Tr |  |  |  |  |
| FFA 16:2 +1O_mz_267.1966_RT_4.28 | Sm, Tr | this study | Copepods | 9-KHME | 5-epHME |
| FFA 16:3 +1O_mz_265.1809_RT_2.99 | Sm, Tr | Lauritano *et al.* 2016; Russo *et al.* 2020; this study | Copepods |  | 9-HHTrE |
| FFA 16:2 +1O_mz_267.1966_RT_3.46 | Sm, Tr, Flowering plants | this study | Copepods | 9-KHME |  |
| FFA 18:4 +1O_mz_291.1967_RT_2.02 |  |  |  |  |  |
| FFA 18:3 +2O_mz_309.2073_RT_7.32 | cyanobacteria, green algae, moss |  |  |  |  |
| FFA 16:4 +2O_mz_279.1602_RT_1.85 | Sm | this study | Copepods |  | 9-HpHTE |

**Table S4.** Location and data of collection for the dissolved meta-lipidomic samples across the DYEatom cruise track

| Station | Depth (m) | Latitude N | Longitude W | Date | MLD |
| --- | --- | --- | --- | --- | --- |
| 1 | 10 | 36°46.864 ' | 121°59.1257 ' | June 27, 2013 | 7 |
| 4 | 2 | 36°27.31 ' | 122°20.32 ' | June 29, 2013 | 16 |
| 4 | 4 | 36°27.31 ' | 122°20.32 ' | June 29, 2013 | 16 |
| 6 | 4 | 36°42.61 ' | 123°29.62 ' | June 30, 2013 | 26 |
| 6 | 35 | 36°42.61 ' | 123°29.62 ' | June 30, 2013 | 26 |
| 7 | 8 | 37°33.42 ' | 123°37.96 ' | July 1, 2013 | 18 |
| 8 | 1 | 38°8.96 ' | 123°5.95 ' | July 2, 2013 | 7 |
| 8 | 4 | 38°8.96 ' | 123°5.95 ' | July 2, 2013 | 7 |
| 9 | 7 | 38°15.89 ' | 123°58.12 ' | July 3, 2013 | 27 |
| 10 | 10 | 37°50.24 ' | 123°6.6 ' | July 4, 2013 | 10 |
| 11 | 2 | 36°46.67 ' | 121°58.83 ' | July 5, 2013 | 11 |

**Table S5.**
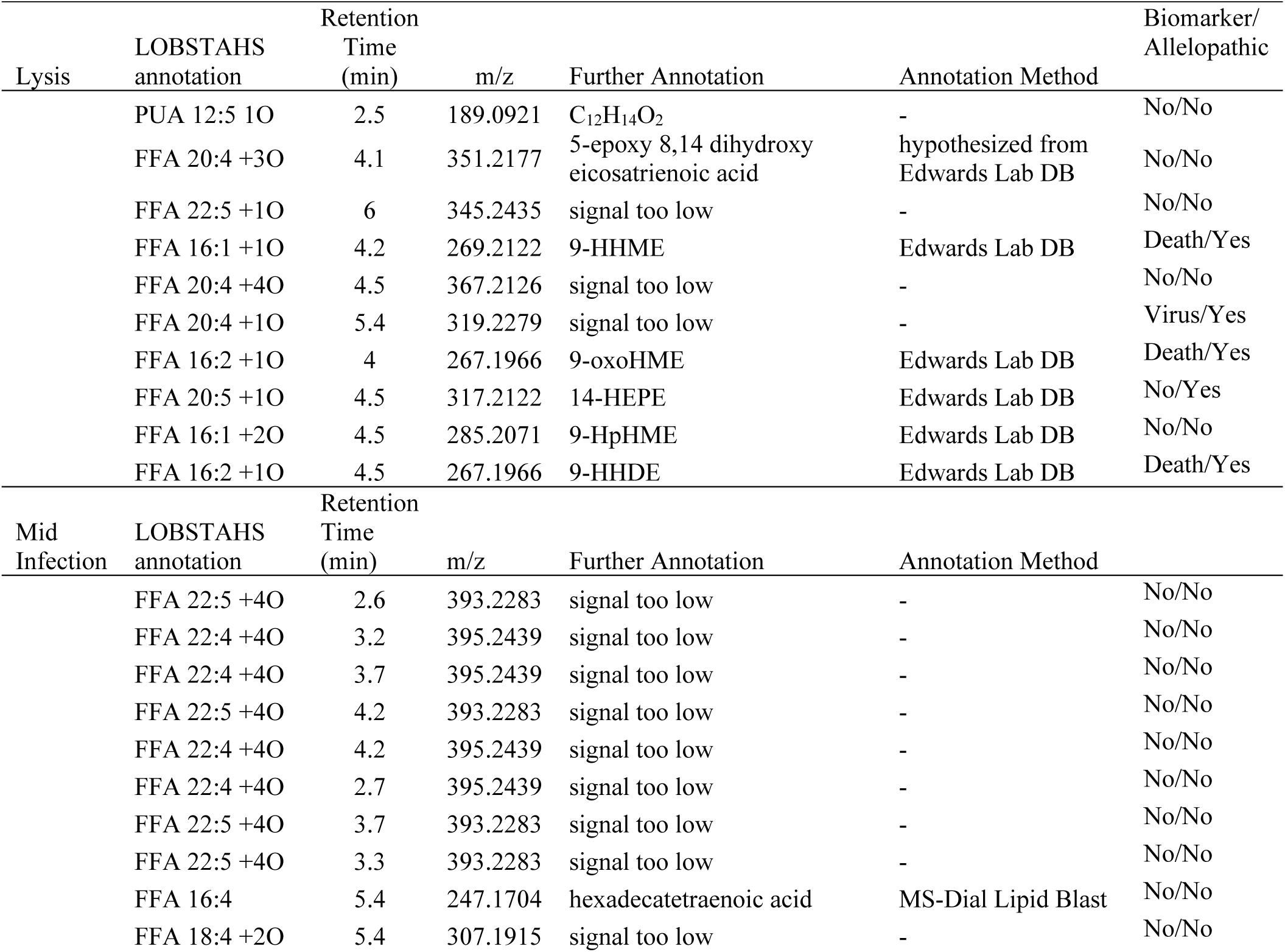
Further annotation of significant compounds identified by sPLS-DA analysis of either lysis or mid infection within the DYEatom cruise dissolved lipidome. LOBSTAHS annotation, peak retention time, the mass to charge ration (m/z) of the feature, and the annotation from considering the ms^2^ fragmentation of samples analyzed using HRAM-MS^n^ analysis outlined in Fig. S3. It is also noted whether each compound was a biomarker observed in the *Chaetoceros* culture experiments in Fig. 3B-C and whether the compound has known allelopathy (based on the studies cited in Table S3).

**Table S6.** Average richness and diversity of the oxylipins and free fatty acids in the dissolved lipidome collected from the surface ocean along the DYEatom cruise. Samples subdivided by the state of diatom infection.

| Non-bloom |  |  |  |
| --- | --- | --- | --- |
| Samples | Richness | Diversity | Sum |
| St6_4m | 165 | -4.03134 | 2x10<sup>8</sup> |
| St6_35m | 164 | -3.97726 | 1.28 x10<sup>8</sup> |
| St9_7m | 156 | -3.92887 | 3.25 x10<sup>8</sup> |
| Avg | 161.6667 | -3.97915 | 2.17 x10<sup>8</sup> |
| stdev | 4.932883 | 0.051261 | 99692081 |
| Post-Lytic |  |  |  |
| Samples | Richness | Diversity | Sum |
| St7_8m | 175 | -3.60247 | 5.27 x10<sup>9</sup> |
| St8_1m | 175 | -3.47831 | 2.31 x10<sup>9</sup> |
| St8_4m | 174 | -3.74992 | 3.74 x10<sup>9</sup> |
| St10_10m | 174 | -3.54632 | 1.4 x10<sup>10</sup> |
| Avg | 174.5 | -3.59426 | 5.48 x10<sup>9</sup> |
| stdev | 0.57735 | 0.115527 | 6 x10<sup>9</sup> |
| Active Infection |  |  |  |
| Samples | Richness | Diversity | Sum |
| St11_2m | 174 | -2.94217 | 2.62 x10<sup>9</sup> |
| St4_2m | 174 | -3.28648 | 1.37 x10<sup>9</sup> |
| St1_5m | 175 | -2.94248 | 2.17 x10<sup>9</sup> |
| St4_9m | 172 | -2.71614 | 1.96 x10<sup>9</sup> |
| Avg | 173.75 | -2.97182 | 2.03 x10<sup>9</sup> |
| stdev | 1.258306 | 0.235316 | 5.18 x10<sup>8</sup> |

**Table S7.** T-test results showing a significant difference between the average diversity of the dissolved lipidome collected during three different states of viral infection. Bold italics values are significant <0.01, bold values <u>are significant <0.05 with a Bonferroni correction.</u>

|  | Richness | Diversity | Sum |
| --- | --- | --- | --- |
| Post-lytic vs. Non-bloom | 0.044609 | ***0.003134*** | 0.177903 |
| Active Infection vs. Post-lytic | 0.336716 | **0.007219** | 0.334421 |
| Active Infection vs. Non-bloom | 0.045554 | ***0.002336*** | **0.004728** |

Supplemental File 1: Dissolved Lipidome Rutgers *Chaetoceros* Viral Infection Experiments

Supplemental File 2: Dissolved Meta-Lipidome DYEatom Cruise.

Supplemental File 3: DYEatome Cruise Lipidome Retention Time Model

